# A Conformation-specific Nanobody Targeting the NMN-activated State of SARM1

**DOI:** 10.1101/2022.03.25.485784

**Authors:** Yun Nan Hou, Yang Cai, Wan Hua Li, Wei Ming He, Zhi Ying Zhao, Wen Jie Zhu, Qiang Wang, Jun Liu, Hon Cheung Lee, Stjepanovic Goran, Hongmin Zhang, Yong Juan Zhao

## Abstract

Upon axonal injury, Sterile alpha (SAM) and Toll/interleukin-1 receptor (TIR) motif containing 1 (SARM1) is activated by nicotinamide mononucleotide (NMN) to deplete NAD and consequently promote the process of axon degeneration (AxD). Currently, only the inactive form of SARM1 in its auto-inhibitory conformation has been resolved. The flexibility of the enzymatically active form of SARM1 has so far precluded its structural determination. To solve the problem, we generated a stabilizing nanobody, Nb-C6, that specifically recognized only the NMN-activated form of SARM1. The conformation specificity was verified by immunoprecipitation and surface plasmon resonance. Fluorescently labeled Nb-C6 could immunostain only the activated SARM1 in cells stimulated with CZ-48, a permeant mimetic of NMN. Expression of Nb-C6 in live cells resulted in stabilization of the active form of the endogenous and exogenous SARM1, producing and elevating cellular levels of cyclic ADP-ribose, a calcium messenger. Cryo-EM of the NMN-activated SARM1 complexed by Nb-C6 showed an octameric structure resembling a “blooming lotus” with the ARM domains bending significantly inward and swinging out together with the TIR domains to form the “petals of the lotus”. Nb-C6 bound to the SAM domain of the activated SARM1 and stabilized its Armadillo repeat motif domain. Analyses using hydrogen-deuterium exchange mass spectrometry (HDX-MS), and cross-linking MS (XL-MS) indicate that the activated SARM1 is highly dynamic and flexible and the neighboring TIRs form dimers via the surface close to one BB loop. The Nanobody is thus a valuable tool for delineating the mechanism of activation of SARM1 in AxD and other cellular processes.

## Introduction

Axon degeneration (AxD) is regarded an early event in various neurodegenerative diseases including amyotrophiclateral sclerosis, multiple sclerosis, and Parkinson’s disease^1^. AxD is a programmed process, regulated by the axonal NAD levels. This is revealed by the *Wld^s^* mice that express a cytosolic NMNAT1, delaying the axonal NAD depletion after injury and reducing AxD dramatically^2,3^. The genetic screening in *Drosophila melanogaster* led to the discovery of the crucial protein, Sterile Alpha (SAM) and Toll/interleukin-1(TIR) Motif–containing 1 (SARM1), ablation of which also significantly delayed AxD^4^.

SARM1 is a multi-domain protein, composing of an N-terminal 27-aa mitochondrial localization signal, an Armadillo repeat motif (ARM) domain, a SAM domain, and a C-terminal TIR domain. Truncation studies suggest that the TIR is the catalytic NADase domain while the ARM serves an auto-inhibitory function^5,6,7^. The enzymatic activity of the auto-inhibited of SARM1 is activated by nicotinamide mononucleotide (NMN) *in vitro* or by CZ-48 in live cells, a permeant mimetic of NMN we previously developed^8^.

In addition to being a substrate of SARM1, NAD has been found to be also an endogenous inhibitor^9,10^ and that the increased NMN/NAD ratio actually regulates SARM1 activation and AxD^11^. These findings established a working model of AxD, in which NMN accumulation and NAD decrease, caused by rapid degradation of NMNAT2 after axon injury^12,13^, activates SARM1. The result is depletion of NAD, leading to AxD.

SARM1 is, in fact, a novel signaling enzyme possessing multi-catalytic functions. In addition to its NADase activity, it also catalyzes the production of two calcium-mobilizing messengers, cyclic ADP-ribose (cADPR) and nicotinic acid adenine dinucleotide phosphate (NAADP)^8,14–16^. In paclitaxel-induced peripheral neuropathy, SARM1 is activated to produce cADPR, which in turn elevates cellular calcium, resulting in AxD^17^. SARM1 is mainly expressed in neurons^18^, but is also found in a variety of cells^8^. It mainly localizes in the outer membrane of mitochondria with the catalytic domain facing the cytosol^19^ and has ready access to the substrate, NAD. All these features, together with its auto-regulatable nature, make SARM1 a potential calcium signaling enzyme with roles not only in AxD^20^ but also potentially other physiological conditions as well.

SARM1 was originally thought to function as a dimer^5,6,8^ until the octameric structure of its SAM domain was shown^21^. Shortly afterward, several structures of SARM1 in its inactive form were solved by cryo-EM technique^9–11,22,23^. In these structures, the ARM domains form the outer periphery of the octameric ring (c.f. the bottom view shown in **Fig. 3e**) with the TIR domains wedged in between, while SAM domains form the inner periphery around the center hole of the ring. The binding site of NMN at the ARM domain has been revealed by the crystal structure of the isolated domain from the *Drosophila* SARM1, dSARM, which occupation can induce a significant conformational change in the entire domain^11^. Despite much effort, the structure of the NMN-activated form of SARM1 has not been determined, largely due to the flexibility of the ARM and TIR domains upon activation.

In this study, we generated and isolated a unique nanobody, Nb-C6, that binds specifically only to the active form of SARM1 and stabilizes it, allowing the first structural determination of the NMN-activated SARM1. Detailed analyses indicate that the binding and activation by NMN induces large conformational changes in the octameric SARM1. Expression of Nb-C6 in live cells activates SARM1 and elevates cellular cADPR through stabilization of the active form. The nanobody thus provide a novel tool to localize and visualize active SARM1in cells as well.

## Results

### Generation and isolation of a nanobody that specifically recognizes to the NMN-activated SARM1

Our approach to determine the structure of the active form of SARM1 was to stabilize it using a nanobody (Nb). We produced the recombinant dN-SARM1, with the N-terminal 27-aa segment truncated and carrying a BC2-tag^24^, in HEK293T cells. The proteins, immunoprecipitated by the beads conjugated with a BC2 nanobody^24^, were mixed with Freund’s adjuvant and immunized an alpaca. After several rounds of immunization, the nanobody library was constructed from the mRNAs of the peripheral lymphocytes. Phage display and clone selection were performed as described in Methods section (**Fig. 1a**). Several nanobodies were cloned, expressed in bacteria and purified with His_6_-tag affinity column using the protocol described previously^25^. One of the nanobodies, Nb-C6, sequence and secondary structure shown in **Fig. 1b**, was selected because it recognized only the NMN-activated SARM1. In the pull-down assay as shown in **Fig. 1c**, Nb-C6 co-precipitated with SARM1 only in the presence of the activator, NMN, while Nb-1053^25^, an irrelevant Nb control, did not bind SARM1, indicating Nb-C6 is specific for the activated conformation.

**Figure 1.**
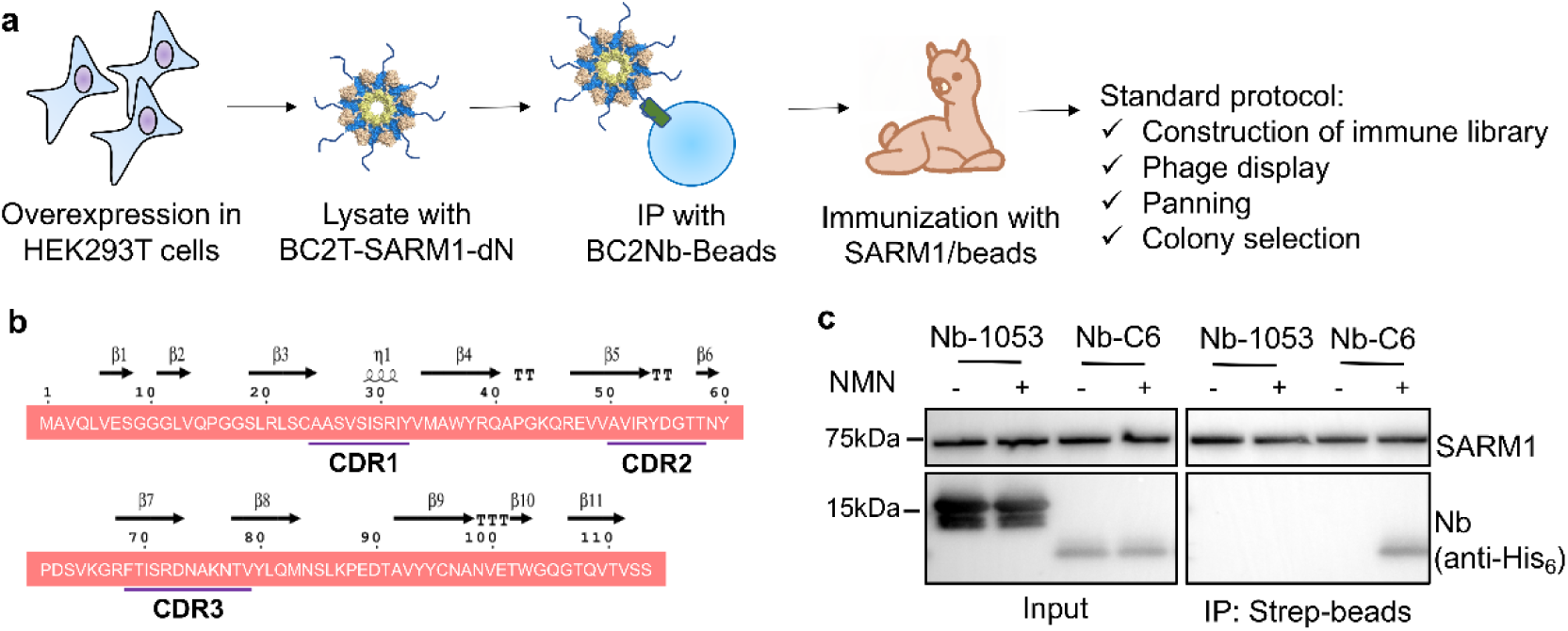
Nanobody C6 binds to the NMN-activated SARM1. (**a**) The process of Nb- C6 development. (**b**) The primary sequence and secondary structure of Nb-C6. (**c**) Nb- C6 binds to the NMN-activated SARM1 determined by the pulldown assay. The cell lysates containing the recombinant SARM1 were incubated with the StrepTactin^TM^ beads, together with 200 ng/mL Nb-C6, or a CD38 nanobody Nb-1053^25^ as a control, in presence or absence of 100 μM NMN. The protein complex was eluted by 2 mM biotin and analyzed by western blots.

### Visualizing and *in vivo* activating SARM1 using Nb-C6

We used surface plasmon resonance (SPR) to confirm that Nb-C6 indeed targeting only the activated form SARM1. As described in the Methods section, purified recombinant SARM1 with a strep-tag was first immobilized on the surface of a SPR chip coated with Strep-Tactin^TM^ XT proteins. An affinity (*K*_D_) of ∼12 nM was measured for the binding of Nb-C6 to the activated complex, SARM1^NMN^ (abbreviation: P^M^ means the protein or domain “P” loaded with the small molecule “M” in the whole text) (**Fig. S1**). The effect of varying the NNM/NAD ratio was assessed next. A running protocol was designed to shift the binding ligand gradually from NAD to NMN and measure the resultant changes in the binding rate of Nb-C6 to the immobilized SARM1. As shown in the right panel of **Fig. 2a**, at high NAD concentration (1.6 mM) the binding of Nb-C6 to SARM1 was very low (blue in scale bar). As the ligand was gradually shifted to NMN, the binding increased more than 80 folds at the highest concentration of NMN (100 µM) (yellow in scale bar). The binding data were fitted with a polynomial model (equation in **Fig. 2a**, left inset) and the binding rate of Nb-C6 to SARM1 was clearly and positively correlated to the concentration of NMN and negatively to that of NAD, with a correlation coefficient R^2^=0.91 (**Fig. 2a**, left inset).

**Figure 2.**
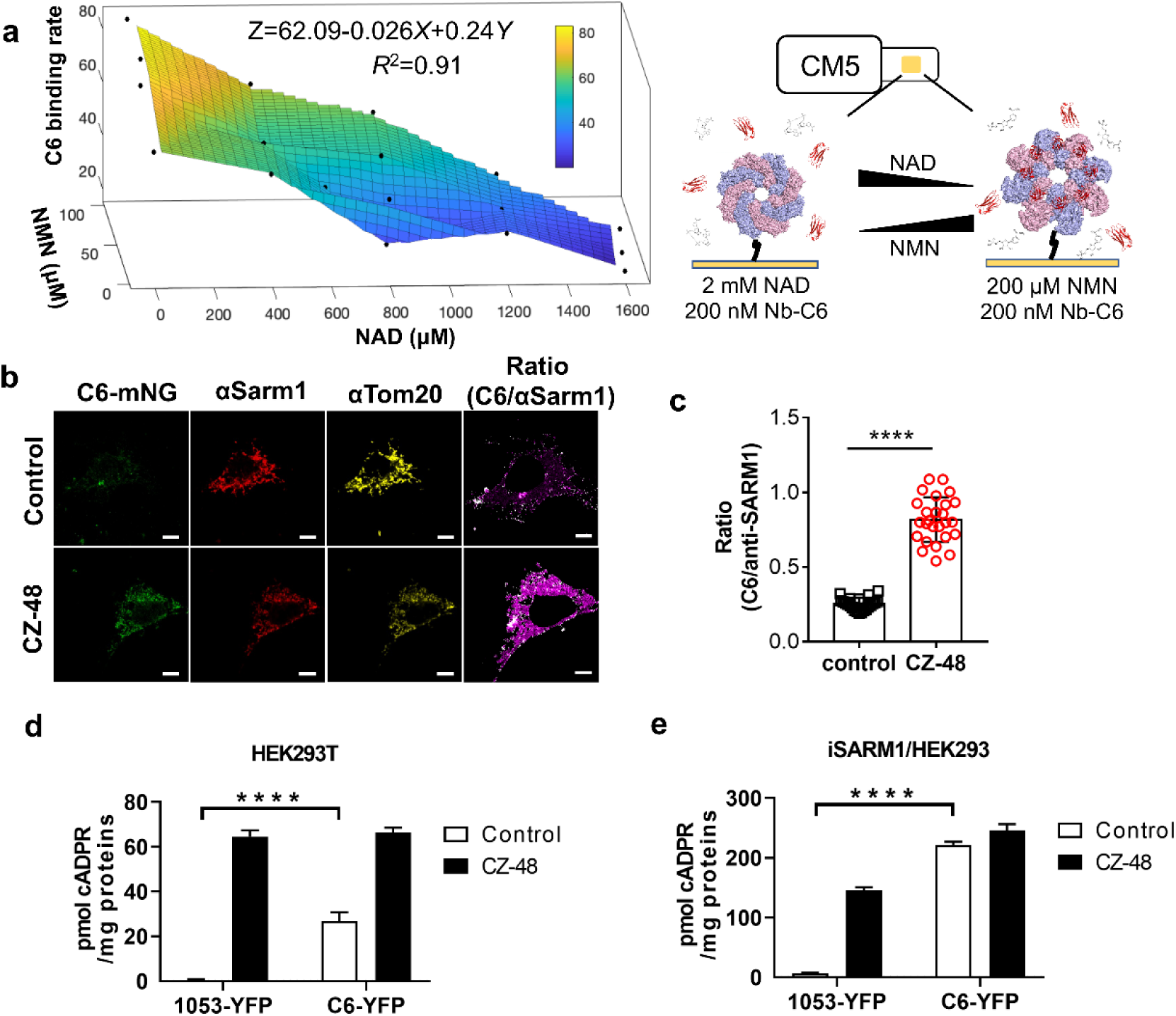
Nb-C6 indicates and facilitates SARM1’s activation *in vitro* and in cells. (**a**) SPR analysis of Nb-C6 binding to the immobilized SARM1 when the mobilizing modulator switching from 2 mM NAD to 200 μM NMN. (**b**) Nb-C6 stains the CZ-48- activated SARM1. HEK293 cells carrying the inducible expression cassette for SARM1 were treated with 1 μg/mL doxycycline for 12 h and then 100 μM CZ-48 for 6h. The cells were fixed and stained with C6-mNeonGreen, anti-SARM1, anti-Tom20 as described in the Methods. Scale bar: 5 μm. (**c**) The ratio of Nb-C6 and anti-SARM1 signals was calculated and plotted. (**d-e**) Nb-C6 elevated the cellular cADPR levels. The HEK293T cells (d) or HEK293 cells carrying the inducible SARM1 expression cassette (e) were transiently transfected with the plasmids encoding the YFP-fusion nanobody, C6 or 1053. The cells were harvested 48 h post-transfection and the cellular cADPR contents were measured by cycling assay and proteins by western blots (Fig. S2e-f). The data are presented as means ± SDs; n = 3.

We next tested if Nb-C6 can detect the activated form of SARM1 in cells. Nb-C6 fused with a fluorescent tag, mNeonGreen (mNG), was produced and used for visualizing the activated SARM1. HEK293 cells exogenously express low levels of SARM1 that can be activated by the permeant NMN mimetic, CZ-48^8^. Positive staining by Nb-C6 was seen in the cells only after CZ-48 treatment, but not in the untreated control cells (C6-mNG, **Fig. 2b**, green signals). For comparison, we generated polyclonal anti-SARM1, raised against the recombinant SARM1 and it could recognize SARM1 in both conditions (αSARM1, **Fig. 2b**, red signals). The Nb-C6 signals were perfectly co-localized with the anti-SARM1 (**Fig. S2a-b**) staining with the coefficient values (PC, M1, and M2)^26^ close to 1. It is believed that SARM1 with its N-terminal localization signal is normally expressed at the outer mitochondrial membrane^8,19^. This is further confirmed here as both the Nb-C6 and anti-SARM1 signals were co-localized with anti-Tom20 (mitochondrial marker; **Fig. 2b**, yellow signals; **Fig. S2c-d**). The Nb- C6 signals were then quantified by normalizing pixel-by-pixel to the anti-SARM1 signal, which represented as the total amounts of SARM1 in the same cell (ratio, **Fig.2b**, last panels). The average of the ratios from around 20 cells (**Fig. 2c**) demonstrated the significant increase of Nb-C6 binding upon CZ-48-induced activation of SARM1. The results confirm the conformational specificity of Nb-C6 and suggest that the activation of the mitochondria-localized SARM1 by NMN in live cells also undergoes conformational changes and exposes the recognition site for Nb-C6.

We reasoned that Nb-C6 itself may be able to promote the activation of SARM1, since its binding site on SARM1 could be exposed randomly at low levels even without activation. The presence of Nb-C6 could then bind to and stabilize the active form. This is the case as shown in **Fig. 2d** and **2e.** Nb-C6, with a soluble tag YFP (C6-YFP), was transfected and overexpressed in two cell types. HEK293T cells endogenously express significant levels of SARM1, which can be activated by CZ-48 treatment^8^, and, when overexpressing C6-YFP (**Fig. S2e**), showed elevated cADPR levels, indicating Nb-C6 alone can activate the endogenous SARM1 (**Fig. 2d**, C6-YFP). No cADPR increase was seen in the control cells overexpressing an irrelevant Nb, 1053-YFP^25^. In another cell line, HEK293 cells expressing inducible SARM1, also showed higher levels of cADPR after transfecting with Nb-C6 as compared to those transfecting with 1053-YFP (**Fig. 2e** and **Fig. S2f**). The increased cADPR levels were actually as high as in CZ-48 treated control cells expressing 1053-YFP, confirming that the expression of Nb-C6 can fully activate exogenous SARM1 as well.

The above results fully validate that Nb-C6 recognizes only the activated form of SARM1 and show that Nb-C6 is a valuable tool for activating endogenous and exogenous SARM1 in live cells.

### Overall cryo-EM structure of the NMN-loaded SARM1 complexed with Nb-C6

We next determined the stabilized structure of the activated SARM1 by cryo-EM. The recombinant SARM1, with the N-terminal 27-aa segment replaced with a Flag/Strep tag, was prepared, and characterized as described previously^23^. The recombinant SARM1 was pre-incubated with NMN and Nb-C6.

Cryo-EM images were collected and processed as detailed in the Methods section (**Fig. S3a-c**). The overall resolution for the structure is around 3.0 Å. Similar to the inactive SARM1, the active form was also octameric. Nb-C6 bound mainly to the SAM domain and also part of the linker between ARM and SAM domains (**Fig. 3c**), stabilizing the whole structure like pillars. The TIR domains were less stable as determined by particle imaging analyses and the most flexible part was the outer ring formed by the ARM domains (ARM1-4) (**Fig. S4**). Using the structural information obtained from crystallography of individual domains, all the residues in SARM1 could be traced except those in the linker between the SAM and TIR domains. The octomer showed a binding ratio of 16:16 between SARM1 protomer and Nb-C6, with two SARM1 octamers bridged by the dimerization of Nb-C6 at their C-termini (**Fig. S3d**). The two octamers in the structure were almost identical, hence we analyzed only one.

The SARM1 octamer complexed with NMN, SARM1^NMN^, was assembled through the oligomerization of the SAM domains (**Fig. 3a-c**, yellow), which was almost identical to that in the previous published SARM1^NAD^ (PDB 7ANW; **Fig. 3d-f**, yellow) with a root mean square deviation (RMSD) of ∼0.62 Å for all Cα atoms in the SAM domain. The overall architecture of SARM1^NMN^ exhibited a “blooming lotus” shape (**Fig. 3a-c**), in sharp contrast to the compact “doughnut” structure of SARM1^NAD^ (**Fig. 3d-f**). The ARM-TIR domains, as a whole, swung out from SAM domain and rotated ∼139°, 73 Å for N-terminal residue E60, as compared to that in SARM1^NAD^ (**Fig. 3g**, blue to green for ARM domain; wheat to magenta for TIR domain). This large conformational change of SARM1 following NMN binding exposed regions recognized by Nb-C6, accounting for its specificity only for the NMN-activated SARM1.

**Figure 3.**
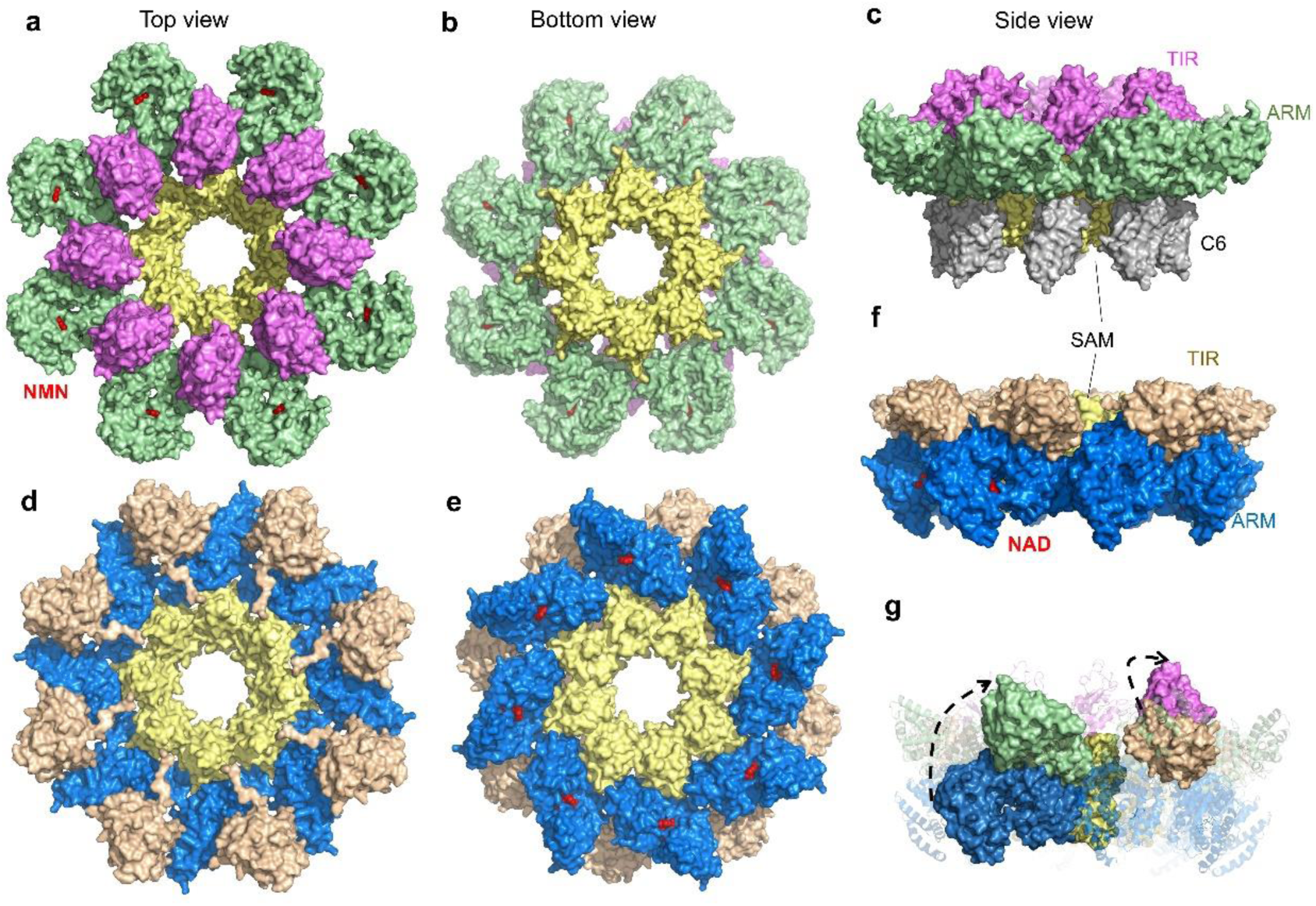
Overall structures of active or inactive SARM1. (**a-c**) Structure of the SARM1^NMN^/Nb-C6 complex. a: top view; b: bottom view; c, side view. SARM1 and Nb- C6 are shown as surface models with ARM domain colored in green, TIR domain in magenta, SAM domain in yellow and Nb-C6 in grey. NMN is shown as sphere in red. Each Nb-C6 binds one SARM1 protomer. (**d-f**) Overall structure of SARM1^NAD^ (PDB 7ANW). d: top view; e: bottom view; f, side view. SARM1 is shown as surface model with ARM domain colored in blue, TIR domain in wheat and SAM domain in yellow. NAD is shown as sphere model in red. (**g**) The side view of superposition of NMN- and NAD-bound SARM1. For clarity, only one protomer in NMN-bound and NAD-bound forms are shown as surface models and other protomers are shown as cartoon, and Nb-C6 in SARM1^NMN^ complex was hidden. SARM1^NMN^ is colored as in panels (a-c) and SARM1^NAD^ is colored as in panels (d-f).

### NMN induces significant structural changes of the ARM domain

NMN bound to the ARM domain, a compact upside down ω-shaped structure formed by eight ARM repeats, each composed of α helices. The binding site was a pocket enclosed by the H3 helix from ARM1, H1 helix from ARM2 and the loop connecting ARM6-7 (named D317 loop hereafter) (**Fig. 4a**). NMN directly interacted with nine residues including W103, R110, E149, Q150, R157, H190, K193, S316, and D317 (**Fig. 4a**, right panel). The pyridine ring of NMN packed against the indole ring of residue W103 forming parallel π-π stack, while the amide group formed hydrogen bonds with the side-chain amide group of Q150 and main-chain carboxyl group of S316, respectively (**Fig. 4a**, right panel). The ribose part of NMN also formed hydrogen bonds with main-chain carboxyl groups of Q150 and E149. The phosphate group formed salt-bridges with the side chains of R110, R157 and K193. The side chain of D317 was close to the pyridine ring of NMN, and formed a hydrogen bond with the amine group in the imidazole ring of H190.

**Figure 4.**
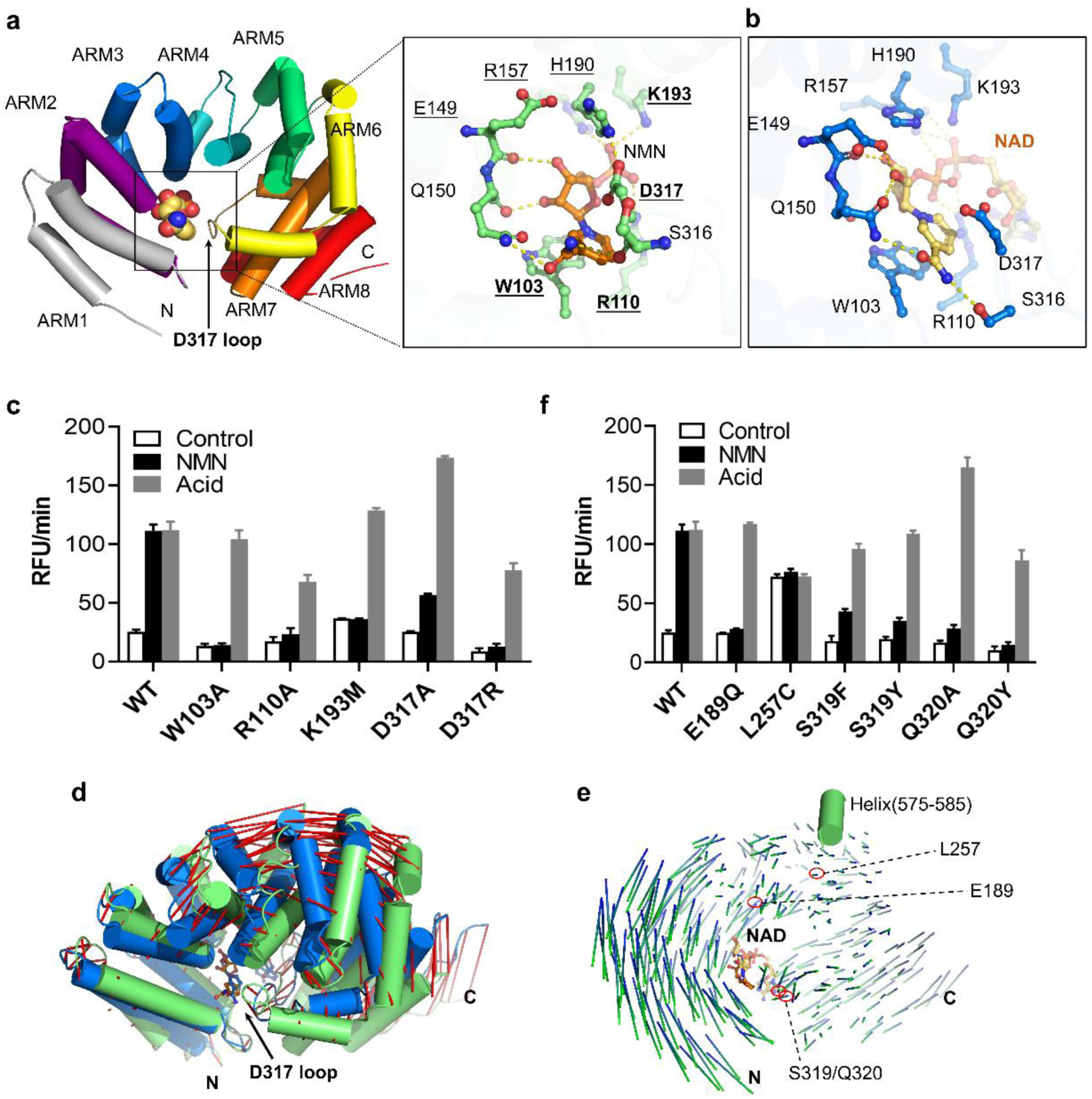
Conformational changes of the ARM domain induced upon NMN binding. (**a**) The structure of ARM^NMN^. ARM domain is shown as cartoon, with separate armadillo motifs in different colors, and NMN molecule is shown as sphere. The D317 loop is indicated with a black arrow. The right panel shows the zoom-in structure of the NMN-binding pocket. NMN and the NMN-interacting residues are shown as ball-and-stick models, with NMN colored with gold carbon atoms and amino acids with green carbon atoms. Hydrogen bonds are shown as yellow dashed lines. The residues mutated in this paper are labeled in bold and the reported mutations with underlines. (**b**) The binding pocket of NAD in ARM domain. SARM1^NAD^ (PDB 7ANW) is superimposed onto SARM1^NMN^ at the same orientation of panel a. Amino acids are shown with blue carbon atoms and NAD with yellow carbon atoms. (**c,f**) The constitutive activity and NMN-responsiveness of the SARM1 mutants. HEK293 cells carrying the inducible expression cassette for the mutants were treated with 1 μg/mL doxycycline for 24 h, the proteins were extracted with PBS containing 4 mM Digitonin and were applied to PC6 assay following the treatment of 100 μM NMN or 0.1 M NaOAc, pH 4.0^23^. c. Mutation of the residues directly interacting with NMN; f. Mutation of the residues involved in conformational change of ARM. (**d-e**) Conformational changes of ARM domain in SARM1^NMN^ (green) compared to SARM1^NAD^ (PDB 7ANW) (blue). Residues 93-104 are aligned in panel (d) and positional shifts of Cα atoms are indicated by red lines. The inward shift of the D317 loop is indicated with a black arrow. In panel (e), residues 575-585 in the TIR domain (green cylinder) are superimposed and the positional shifts of Cα atoms in the ARM domain are indicated by blue-green lines.

The NMN-binding site, in fact, was the same as that for NAD as observed in the inactive SARM1^NAD^ (**Fig. 4b**), accounting for the two ligands being competitive^11^. However, the amide moiety in the nicotinamide group of NAD oriented parallel and formed an H-bond with the side-chain hydroxyl of S316 instead of the main-chain carboxyl in NMN-bound ARM domain. The additional phosphate, ribose and adenosine groups in NAD formed interactions with the D317 loop and pushed it outward ∼5 Å, diminishing the H-bond between D317 and H190 observed in ARM^NMN^. These structural differences can account for NMN being an activator while NAD, an inhibitor. Mutational studies confirm the functional importance of the NMN/NAD binding site.

HEK293 cells were transfected and expressed SARM1 with mutations in residues, W103, R110, K193 and D317 (bold in **Fig. 4a**), and tested for the enzymatic activity and responsiveness to NMN activation, using our recently-developed protocols^23^. As expected, all the mutants became irresponsive to NMN (**Fig. 4c**, black bars vs. white bars; expression level in **Fig. S5a**), while all still could be activated by acid (**Fig. 4c**, gray bars). We have recently shown that acid activates SARM1 by breaking salt bridges between ARM and TIR, resulting in full expression of its enzymatic activities^23^. The results support that these residues are critical to NMN binding. It is also consistent with the previous results showing that similar mutations (underlined in **Fig. 4a**) can eliminate the responsiveness to nicotinamide ribose-induced NMN accumulation in neurons^11^.

Next, we investigated the NMN-induced conformational change of ARM domain by aligning ARM^NMN^ (**Fig. 4d-e**, green) and ARM^NAD^ (blue). Large conformational changes were clearly visible after aligning the H3 helix of ARM1, residues 93-104, as indicated by the red-line shift in **Fig. 4d**. ARM^NMN^ showed significant inward bending compared to ARM^NAD^ with the C-terminal ARM7-8 motifs moved much closer toward the N-terminus of ARM domain, pushing the D317 loop inserting more deeper into the NMN binding pocket. Similar large changes were seen by superimposing residues 575-585 in the ARM:TIR interface (**Fig. 4e**, green cylinder). The length of the lines in **Fig. 4e** indicates the movement distance of individual residues, while the color blue (ARM^NAD^) to green (ARM^NMN^) indicates inward direction of the movement. The large conformational change is also evidenced from the large RMSD (∼2.7 Å) yielded from the superposition between ARM^NMN^ and ARM^NAD^ of hSARM1, which is mainly attributed to the shift of helices in the convex side and also the inward movement of ARM7-8 motifs (**Fig. 4d**).

It is likely that the inward bending of the ARM domain induced by NMN is necessary for the enzymatic activation of SARM1. To test this, we mutated several residues that potentially could contribute to the bending, including E189, L257, S319 and Q320 (locations indicated in **Fig. 4e**). The choice of these residues was suggested by structural modeling indicating that replacing S319 or Q320 with a tyrosine could elicit strong hydrophobic interactions with residues in the H3 helix of ARM1 and the H1 helix of ARM2 and hence could interfere the inward movement of the D317 loop upon NMN binding (**Fig. S5b**). Consistently, enzymatic activity of the mutants showed that the NMN-induced activation was abolished in most of these mutants as shown in **Fig. 4f** and **Fig. S5c.** An exception was L257C, which was constitutively active, with its basal activity already being similar to that activated by NMN or acid. Structural modeling in **Fig. S5d** indicates that the cysteine side chain of L257C could reduce the hydrophobic interaction and might form disulfide bond with C215 as well, promoting the inward bending of ARM4-5 and rendering the mutant constitutively active. These results support that the inward bending of ARM domain is necessary for NMN-induced SARM1 activation.

### NMN-induced release of the ARM domain from SAM

Next, we investigated how NMN-induced bending of ARM leads to its dissociation from SAM domain. The ARM domain of each protomer interacts with both its own SAM (intra-chain) and the adjacent SAM (inter-chain) as well. **Fig. 5a** shows the involved interfaces. When complexed with NAD, two key residues in ARM were critical for the intra-chain ARM:SAM association (**Fig. 5a**, upper left panels). Residue R376 formed salt-bridges with E469 in SAM, while Y380 was inserted into a hydrophobic pocket formed by residues I405, W420, L443, L470 and F476. Especially, the π-π stack between Y380 and F476 was pivotal in maintaining the ARM-SAM association. Mutating F476 to cysteine destroyed the stacking and released the ARM:SAM association, rendering the mutant constitutively active (**Fig. 5b**). Likewise, the mutants R376A, E469A and Y380A (underlined in **Fig. 5a**) also had increased basal activities^27^, supporting that the intra-chain ARM:SAM interactions are critical for maintaining the auto-inhibitory state. In the inter-chain ARM:SAM’ interface (**Fig. 5a**, upper right panel), several H-bonds and salt-bridges were observed. These inter-chain association, however, seemed not contributing to auto-inhibition because all the tested mutations (italic in **Fig. 5b**, upper right panel) behaved like wildtype^27^.

**Figure 5.**
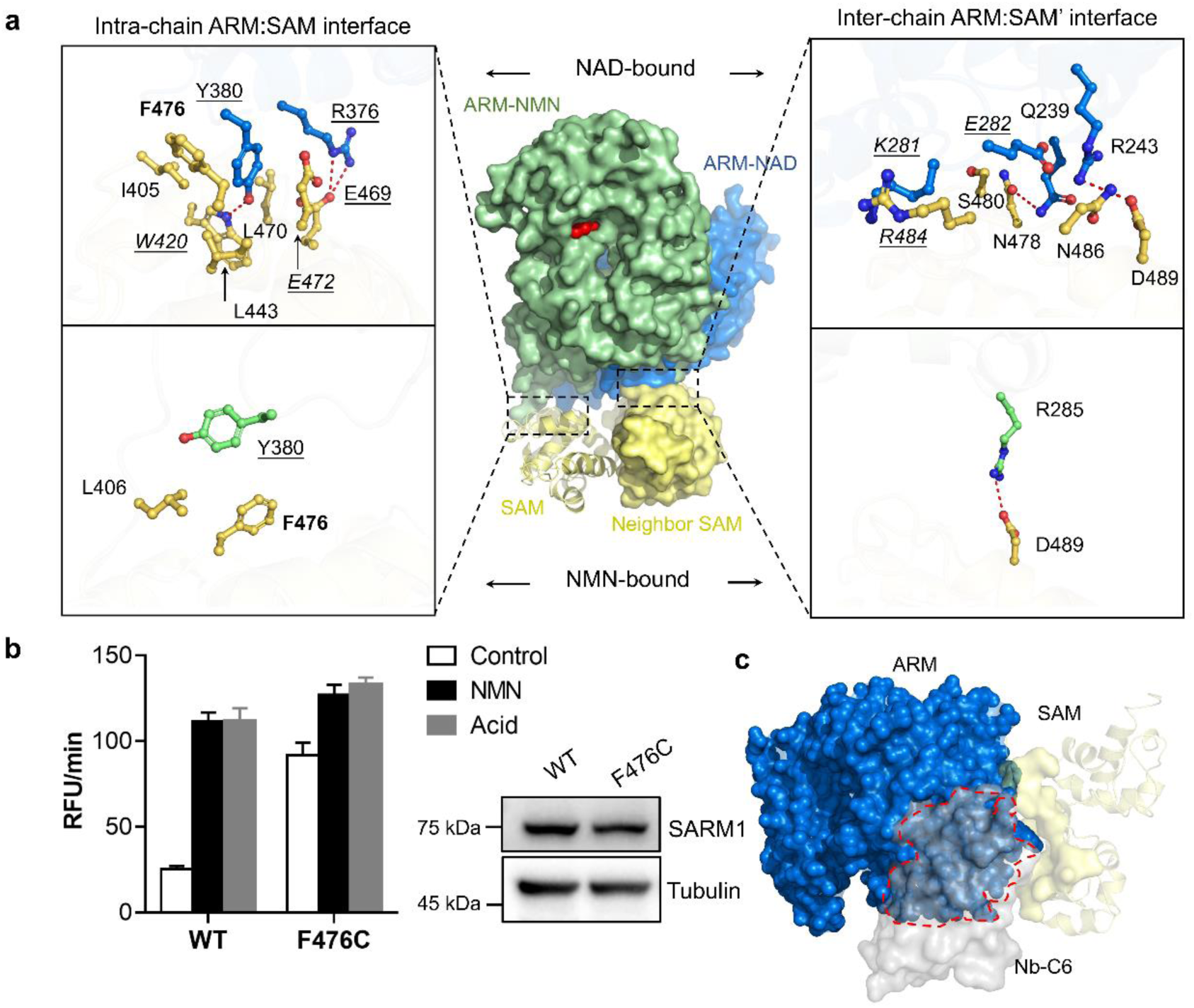
Releasing ARM from SAM domain allows Nb-C6 binding. (**a**) Superposition of SARM1^NMN^ onto SARM1^NAD^ (PDB 7CM6) via the SAM domains. Left panels: intra-chain ARM-SAM interfaces; Right panels: inter-chain ARM-SAM’ interfaces; Upper panels: NAD-bound; lower panels: NMN-bound. The ARM and SAM domains are shown in green and gold in SARM1^NMN^, respectively; and they are colored in blue and orange in SARM1^NAD^, respectively. Key interaction residues at the interfaces are shown in ball-and-stick models and labeled. The H-bonds and salt- bridge are shown as red dash lines. The residues mutated in this paper are labeled in bold and the reported mutations with underlines; the ones behave similar to wild-type labeled in italic. (**b**) The activity and NMN-responsiveness of the SARM1 mutants measured as Fig. 4c. (**c**) Superposition of SARM1^NMN^ onto SARM1^NAD^ (PDB 7CM6) via the SAM domains. The proteins are shown as surface models with SAM domain in yellow, nanobody in grey and ARM domain of SARM1^NAD^ in blue. For clarity, ARM domain of SARM1^NMN^ is not shown. The overlapping area was highlighted by a red dash line.

When complexed with NMN, both the intra- and inter-chain ARM-SAM associations were diminished, with Y380 flipping away from F476 in the intra-chain SAM and only one salt-bridge (R285-D489) remained in the inter-chain interface (**Fig. 5a**, lower panel). The binding of NMN to ARM induced large conformational changes in the domain, comprising of an inward shift of ARM7-8 repeats toward the NMN-binding pocket and the rotation of the whole ARM domain (**Fig. S6a**). As a result, the distances increased between all the interacting residues observed in the SARM1^NAD^ complex, rendering their interactions ineffective. The diminished interactions release SAM from ARM and exposed the site for Nb-C6 to bind. The nanobody mainly bound to the SAM domain with a less extent to the ARM-SAM linker (**Fig. S6b**). The binding interface between Nb-C6 and SAM domain was largely overlapped with the intra-chain ARM:SAM interface (**Fig. 5c**), accounting for the fact that Nb-C6 does not recognize inactive SARM1 as the binding site is obscured by ARM.

### NMN induces conformational changes in the TIR domains

Upon binding of NMN to SARM1, the ARM and TIR domains swung out as a whole with minimal changes in the interface between them (**Fig. S7a**). The movement and the accompanying bending of the ARM domain described above, however, did result in shifting away of the R216 residue in the intra-chain ARM domain (**Fig. S7a,** black arrow) from E689 in the TIR domain, breaking the salt bridge between them. The linkage was believed to be also essential for its auto-inhibition^23,27^.

Superimposing TIR from SARM1^NMN^ with that from SARM1^NAD^ showed that the overall structures were almost identical including the BB loop (**Fig. S7b**, magenta/blue loops), which is considered to be important in determining the enzymatic activity state^6,28^. The conformation of BB loop in both complexes appeared to be constrained by the linker between SAM and TIR (**Fig. S7b**, in sphere mode) and might hinder the entrance of substrate into the catalytic pocket. This may not be of much concern, however, since the TIR domain exhibits high degree of flexibility in solution as shown by HDX-MS and XL-MS experiments described below.

SARM1 was pre-incubated with the excess of either NAD or NMN and followed by HDX labeling for various periods, starting with 10 s, which is sufficient to deuterate fully solvated amides^29^. By using a combination of pepsin and guanidinium denaturant, peptide coverage was obtained for the majority of the homo-octameric SARM1 complex. Altogether, 403 peptides common to both NAD- and NMN-liganded SARM1 were chosen for data analysis. The HDX data are largely consistent with the α helices in ARM, SAM and TIR domains, as observed in the cryo-EM structures. These secondary structure regions incorporated deuterons to a much lower degree than the interconnecting loop regions. Amides involved in stable hydrogen bonds, such as in α- helices and β-sheet secondary structure, exchange several orders of magnitude slower than exposed hydrogens (**Fig. 6a** and **Fig. S8**). Interestingly, the peptide isotopic peaks corresponding to helix α13 (exemplified by peptide spanning residues H236-F259) in the ARM domain formed a clear bimodal distribution in SARM1. A bimodal distribution could result from EX1 type HDX kinetics, in which two distinct protein conformations with different solvent accessibility interconvert during the HDX incubation period^30,31^.The hydrogen exchange observed in these peptides can be described by EX1 kinetic regime where the unfolding rate is much faster than the refolding rate. Under these conditions, all the amide hydrogens exchange with deuterium in the open state before refolding occurs, resulting in two distinct mass envelopes. The lower mass envelope represents the closed state (**Fig. 6b**, blue shade), and the higher mass envelope represents the open state (**Fig. 6b**, yellow shade). The two molecular mass envelopes were localized to the ARM-TIR interaction interface of the ARM domain (helix α13) (**Fig. 6c**). This behavior was evident at 10 s to 30 min of exchange. In SARM1, the less accessible molecular mass envelope gradually converted into a more accessible state (**Fig. 6b**). Importantly, preincubation with NAD significantly stabilized the less accessible state and slowed down the interconversion between the two states when compared to SARM1^NMN^ (**Fig. 6b, bottom plot**). ARM helix α13 is responsible for direct interaction and recruitment of TIR domain to the periphery of SARM1 homo-octameric ring. Furthermore, NAD-bound state showed reduction in deuteration levels in peptide spanning residues 353-389 and 483-498 located at ARM-SAM interaction interface (**Fig. 6a**). These changes were consistent with a major stabilization effect on the SARM1 dynamics when NAD was present, and support the model where binding of NAD stabilizes interaction between TIR, ARM and SAM domains. In contrast, SARM1^NMN^ complex formed a more dynamic structure, where the TIR domain was dissociated from the ARM domain resulting in faster conversion of helix α13 to the open state.

**Figure 6.**
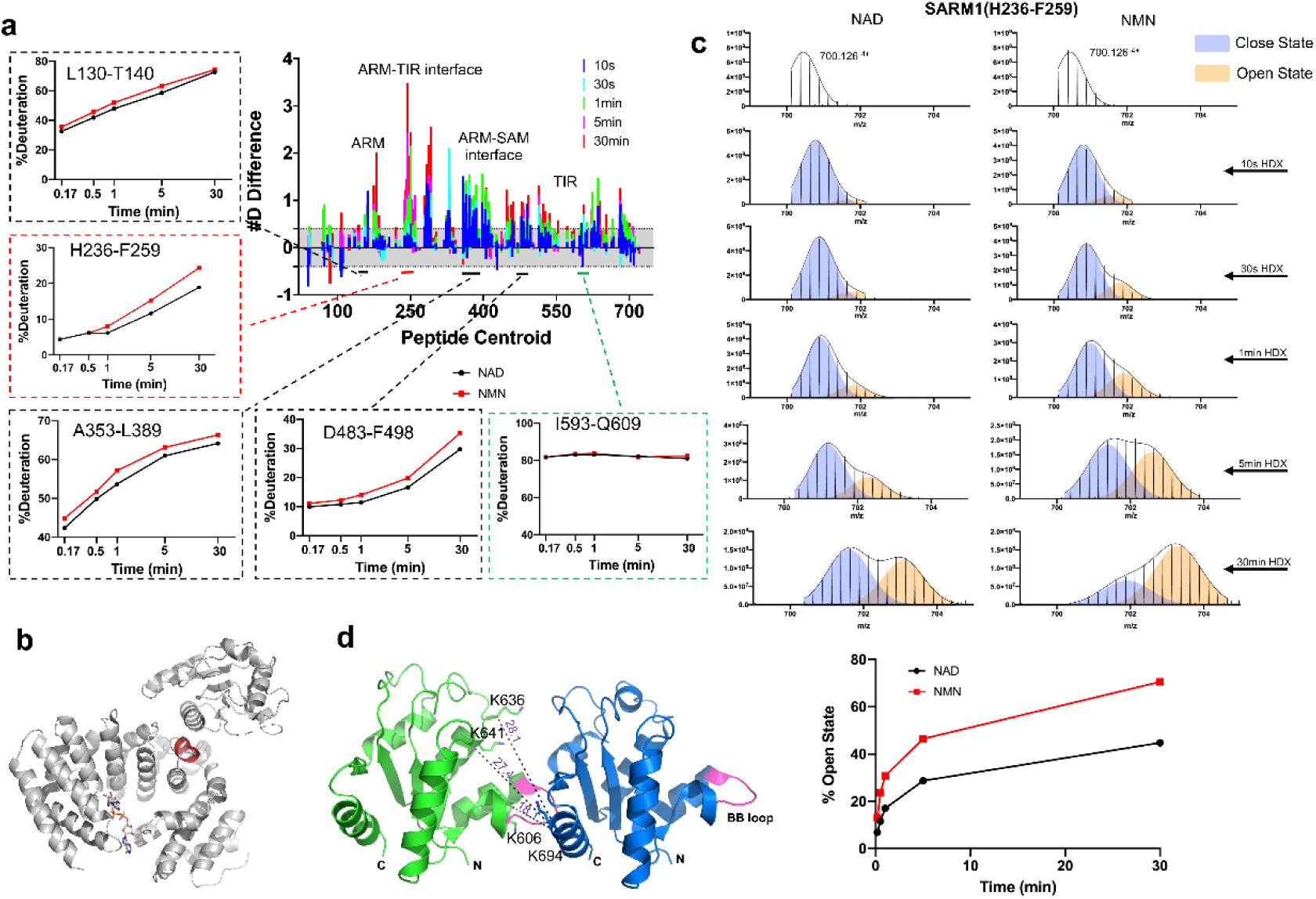
HDX-MS and XL-MS reveal the conformational dynamics of NMN- and NAD-bound SARM1 octamers in solution. (**a**) Bar plot representing differences in deuterium uptake between NMN- and NAD-bound SARM1, with each bar representing the central residue of an individual peptide (ordered from N- to C- terminus). The Y- axis represents the total difference in number deuterium uptake for a given peptide, between NMN and NAD bound SARM1 at each time point. Positive bars indicate increase in exchange of the corresponding peptide in the presence of NMN and reveal that several distinct regions are affected by ligand binding. Light grey shades represent the threshold above which deuterium uptake differences were considered significant (>0.4 Da). Time course of HDX incorporation in presence of NAD or NMN for several representative peptides are shown. (**b**) Bimodal isotopic envelopes for the SARM1 helix α13 (peptide spanning residues 236-259). Relative amount in the closed and open state in presence of NAD and NMN as seen by HDX. Deuteration time points are indicated. Open to close transition kinetics for SARM1 helix α13 is calculated by fit of two gaussians. Next, the fitted peak area parameter was used to calculate the relative amount of open state by taking the ratio of high and the sum of high and low mass subpopulation. (**c**) Structure of the NAD bound SARM1 showing ARM-TIR domain interface with the ARM helix α13 indicated in red. NAD ligand is shown as stick model. (**d**) Mapping the XLs on the TIR dimer (extracted from PDB 7NAK), with a BB loop (colored in pink) as the interface, in the polymer (PDB 7NAK). The distances were labeled.

We next use crosslinking–mass spectrometry (XL-MS) to compare the conformational differences between SARM1 complexed with NMN and NAD. We used the MS-cleavable cross-linker DSBU (disuccinimidyl dibutyric urea), that connects primary amine groups of lysine residues within Cα-Cα distances up to ∼30 Å^32^. Three different SARM1 concentrations (1.25µM, 2.5 µM and 5 µM) were used, at which the protein forms multimers in solution. SARM1 was preincubated with saturating concentrations of NAD or NMN prior to crosslinking. We then used the information derived from XL-MS and cryo-EM for modeling of SARM1 regions of interaction, based on a maximum 30Å Cα–Cα distance constraints between two crosslinked lysine residues. Since SARM1 forms multimers, XL-MS approach used in this study is not capable of differentiating intramolecular from intermolecular links. Interpretation of links as intra- or intermolecular was therefore aided by high resolution cryo-EM structures.

For SARM1 complexed with NAD, a total of 57 crosslinks (XLs) were detected (**Supplementary Table 1**, **Fig. S9a**). Half of the intra- and inter-molecular XLs identified (29 of 57) were consistent with the domain structures and complex architecture observed in the cryo-EM structure of full-length SARM1 (PDB 7CM6) (**Fig. S9b**). The other 29 were unsatisfied cross-link of residues with distances >30 Å (**Fig. S9b**), indicating extensive conformational changes of the domains when in solution. Similarly, large number of unsatisfactory XLs (40 of 62) were observed in the SARM1^NMN^ cryo-EM structure obtained in this study (**Supplementary Table 2**, **Fig. S9c-d**). These XLs were mapping to ARM and TIR domains, indicating significant flexibility within SARM1 complex.

Interestingly, four cross-links (K636-K694, K641-K694, K556-K556, and K606- K694) were readily identified in the NMN treated sample. Three XLs (K636-K694, K641-K694, and K606-K694) involve lysine residues in regions of defined secondary structure located on opposite faces of the TIR domain. These residues were unlikely to form intramolecular XLs since the vectors for these XLs passed directly through the TIR domain. We believe these XLs arose as a consequence of intermolecular TIR-TIR domain interactions. This was further supported by XL dimer (K556-K556) mapping to the loop connecting SAM and TIR domains. We mapped these three XLs on the newly-published cryo-EM structure of a TIR polymer (PDB 7NAK)^33^. The distances of all three residue-pairs were within 30 Å in the dimer with BB loop as the interface (**Fig. 6d**), indicating this might be the predominant interaction. Our data did not provide evidence for the larger oligomeric state of the TIR domains, at least in our experimental conditions.

Taken together, the HDX-MS and XL-MS data suggest that the activated SARM1 in solution is highly dynamic and flexible, making it un-observable by cryo-EM. The Nb- C6 stabilized structure thus may represent an intermediate toward this final state of activation. As depicted in **Fig. 7**, in this final state, the TIR is completely released from ARM and directly interacts with the neighboring TIR, allowing the crosslinking observed with XL-MS.

**Figure 7.**
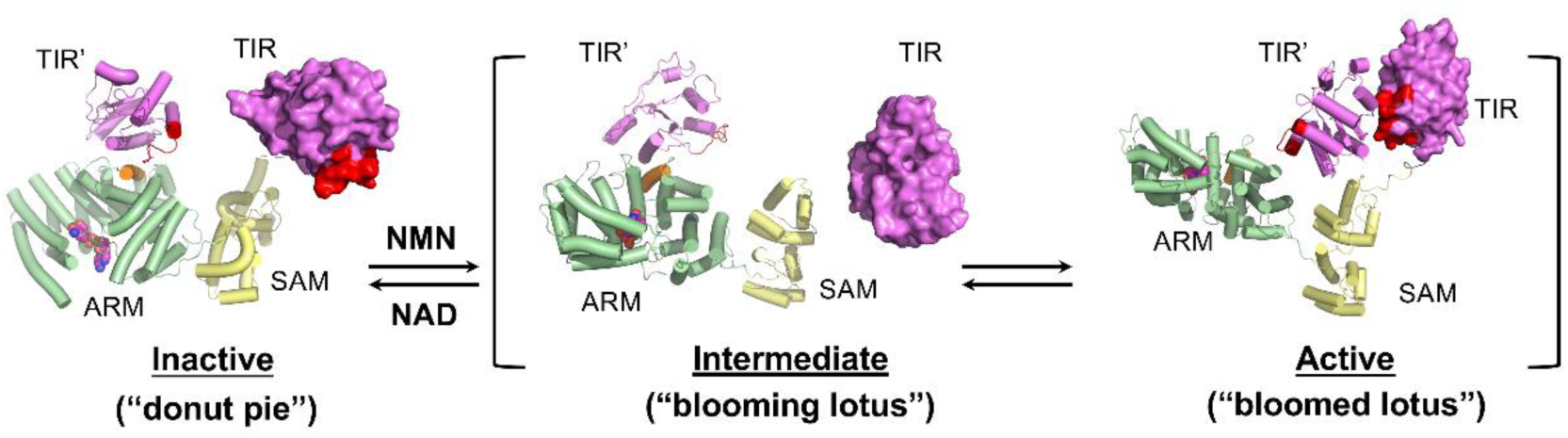
The transition between the inactive and active states of SARM1. SARM1 was shown with the ARM domain in green, SAM domain in yellow and TIR domain in magenta. TIR’ is the domain from a neighboring SARM1 protomer. The 13 helix (residues 249-259) in the ARM domain was colored in orange. BB loop in the TIR domain was colored in red.

## Discussion

SARM1 has attracted substantial interests because of its role in AxD^4^. The finding that it is an NADase^7^ has led to the proposal that its activation after axonal injury can deplete cellular NAD and cause AxD. Recent results show that SARM1 is not just a degradative NADase, but an auto-regulated signaling enzyme responsible for producing two calcium mobilizing messengers, cADPR and NAADP^8,23^. For example, in paclitaxel-induced peripheral neuropathy, SARM1 is activated to produce cADPR, resulting in calcium increase and AxD^17^. We and others have shown that the enzyme activities of SARM1 are specifically activated by NMN^8,11^. Understanding the mechanism of its activation is clearly important and a structural approach is the most direct.

Combining results obtained using cryo-EM and crystallography of various domains have produced a high-resolution structure of the octameric SARM1 in the inactive state^9–11,22,23^. In this study, we showed that the activated state exhibits considerable flexibility as revealed by increased rate of isotopic incorporation, particularly in the helix α13 of ARM (**Fig. 6c**), This flexibility had prevented the resolution of its structure. We circumvent this difficulty by generating and isolating Nb-C6 that binds only to and stabilizes the activated state of SARM1, allowing the determination of its high-resolution structure by cryo-EM.

The activated SARM1 is also an octamer assembled through the SAM domains. In contrast to the donut-shape of the inactive SARM1 complexed with NAD (**Fig. 3d-f**; **Fig. 7**, left panel), the NMN-activated form resembles a “blooming lotus” shape, with ARM and TIR extended out to form the “petals” (**Fig. 3a-c**; **Fig. 7**, middle panel). The nanobody binds mainly to the SAM and partially with the ARM domains, covering essentially the hinge between SAM and ARM and stabilizing the whole structure. The NMN-loaded ARM domain exhibits a significant inward bending, as compared to ARM^NAD^. The NMN-induced conformational change of ARM leads to its releases from SAM, and consequently, the activation of SARM1.

Four lines of evidence indicate that the SARM1 structure presented in this study indeed represents the activated state. Firstly, Nb-C6 immunoprecipitates SARM1 only when it is activated by NMN (**Fig. 1**). Secondly, SPR documents that purified SARM1 can bind Nb-C6 only when it is complexed with NMN (**Fig. 2**). Thirdly, immuno- fluorescence staining by Nb-C6 is observed only when cells are activated by the permeant NMN mimetic, CZ-48 (**Fig. 2**). Lastly and most importantly is that Nb-C6 expresses in live cells can activate endogenous or induced SARM1 to produce cADPR (**Fig. 2**).

Furthermore, structural features resolved in the activated SARM1 octamer are consistent to those previously obtained in individual domains. In particular, the NMN binding site resolved in the activated SARM1 (**Fig. 4**) is essentially the same as that observed in the isolated ARM-NMN complex from *Drosophila* SARM1, dSARM^11^. We aligned and analyzed these two structures in **Fig. S10**.

The active structure stabilized by Nb-C6 is rigid enough for resolution by cryo-EM. It clearly does not represent the highly flexible state in solution we show in this study (**Fig. 6**). It is thus only a single sub-state or intermediate toward the final and fully activated SARM1, which is flexible and cannot be resolved by cryo-EM or crystallography. In this intermediate, the TIR domains still sit upon the ARM domains and remain separated from the neighboring TIR, which association is required for catalytic activities.

Proceeding to the final active state, the TIR is released from ARM and can be chemically crosslinked to its neighboring TIR of the active octamer, as we shown using XL-MS (**Fig. S9**). This is consistent with the results showing that isolated TIR domains must dimerize before they become enzymatically active^6^. The interface of the TIR dimers mainly composed of weak interactions, allowing the TIR domains in the dimers to be very dynamic, inter-converting from complexing with ARM in the intermediate state (**Fig 3** and **S7**) to dimerizing with the neighboring TIR in the fully active state (**Fig. 7**, right panel). The active octamer is thus more flexible and dynamic, resembling a “bloomed lotus”.

The limitations of crystallography and cryo-EM for structural determination are well known. Packing protein molecules in a crystal can introduce distortions. Cryo-EM has fewer packing changes but is still limited by flexibility of the molecules. In this study we used HDX- and XL- MS to assess the dynamics of the fully active SARM1 in solution. This combination of approaches has provided better understand of the activation mechanism of this novel auto-inhibited signaling molecule.

Summarizing, Nb-C6 not only has provided important understanding in the structural changes during the activation process, but also allows imaging and localization of the active SARM1 in cells. Nb-C6 thus supplements well with the permeant probe, PC6, we previously developed for visualizing SARM1 activity in live cells^23^. Furthermore, that transfection and expression of Nb-C6 can activate endogenous SARM1 provides a mean to manipulate SARM1 in live cells in addition to the permeant activator CZ-48 we have synthesized^8^.

## Materials and Methods

### Bacteria and cell lines

OverExpress^TM^ *E. coli* C43(DE3) obtained from Weidi Biotechnology (Shanghai) was used to express the recombinant proteins.

Expi293F cells, obtained from Sino Biological Inc., were cultured in serum-free SMM-293TII media (Sino Biological) at 37°C in a humidified, 5% CO_2_ atmosphere incubator, shaking at 150 rpm. HEK293 and HEK293T cells obtained from ATCC were cultured at 37 °C with 5% CO_2_ in DMEM supplemented with 10% FBS and 1% penicillin/streptomycin.

### Reagents

NAD, NMN, Digitonin, Poly-L-lysine, KH_2_PO_4_, NH_4_HCO_3_, chloroacetamide, polyethylenimine (PEI), and urea were purchased from Sigma-Aldrich. DMEM, Trypsin-EDTA, penicillin/streptomycin solution, Lipofectamine 2000, formic acid, and acetonitrile were purchased from Thermo Fisher. FBS was obtained from PAN Bitotech. SMM-293TII media was obtained from Sino Biological. Ni-Excel column, HiTrap Q column, CM5 sensor were obtained from Cytiva. Strep-Tactin resin, Strep-Tactin XT SPR kit were obtained from IBA. General chemicals were purchased from Aladdin, Macklin, or Sangon Biotech (Shanghai).

### Immunization and screening for Nanobodies (Nbs)

The immunization and library construction were conducted in Shenzhen KangTi Life Technology Co., Ltd (AlpaLife) under approved protocols. Briefly, a healthy male alpaca was immunized with the recombinant dN-SARM1, full-length SARM1 with the N-terminal 27-aa segment replaced by a BC2T-peptide, which was expressed in HEK293T cells^8^ and immunoprecipitated by BC2Nb-conjugated beads^24^. The antigen on beads, mixed with Freund’s adjuvant, was injected subcutaneously for 6 times during 3 months. A VHH library was constructed from the mRNAs of the immunized peripheral lymphocytes. Library screening was performed by the routine protocol of phage display, panning and colony ELISA^34^ using the purified dtSARM1, the full-length SARM1 with the N-terminal 27-aa segment replaced by a twin-Strep and Flag-tag, so called double tag (dt), as the immobilized antigen, which was also used in the cryo-EM study. The phagemid vectors from the positive colonies were sequenced and analyzed by Sangon Biotech (Shanghai).

### Preparation of Nbs or Nb-fusion proteins

The DNA sequence of Nb-C6, or Nb-1053 against CD38 as a negative control^25^, was amplified from the phagemid by PCR and subcloned to pET22b or pXG- mNeonGreen (from Prof. Chu Jun in SIAT/CAS, Shenzhen). The strategies for expression and purification of the recombinant proteins were described as previously^25^.

Briefly, OverExpress^TM^ *E.coli* C43(DE3) was transformed with the expression vector and grown in LB media containing kanamycin or ampicillin at 37℃ until the culture reached at OD_600_ of 1.0. Protein expression was induced by 0.5 mM IPTG at 16℃ for 20 h. Proteins were extracted by French press (ATS) at 800 psi for 3 rounds in lysis buffer (50 mM Tris pH 8.0, 500 mM NaCl) containing 1 mM PMSF and cell lysates were clarified by centrifugation at 20,000 rpm for 45 min at 4℃. The proteins were purified with Ni-excel resin (Cytiva) and HiTrap Q (Cytiva), sequentially, and concentrated by 10kDa Amicon (Millipore).

### Preparation of dtSARM1

The DNA sequence encoding dtSARM1, the truncated form of SARM1 without the N-terminal mitochondrial signal, fused with a twin-Strep-tag and a Flag-tag attached to the N-terminus, were subcloned into pENTR1A-GFP-N2 (#19364, Addgene), then transferred to pLenti-puro (#17452, Addgene) by Gateway^TM^ system (Thermo Fisher). The lentiviral particles were prepared by transfecting HEK293T cells with pLenti- dtSARM1, psPAX2 (#12260, Addgene), and pMD2.G (#12259, Addgene). The Expi293F cell line stably expressing dtSARM1 was established by the lentiviral infection and puromycin selection.

Then, the cells were cultured in a large volume and harvested. The dtSARM1 was released from the cytoplasm by incubating with ice-cold PBS containing 200 μM Digitonin and protease inhibitor (Roche).The cleared lysate was incubated with StrepTactin^TM^ resin (IBA) overnight at 4℃. After washing with ice-cold Buffer W (100 mM Tris pH8.0, 150 mM NaCl, 3 mM EDTA), the proteins were eluted by 2 mM biotin in buffer W and concentrated with 100kDa Amicon (Millipore). The concentration of dtSARM1 was determined by BCA assay (Thermo Fisher).

### Immunoprecipitation and western blots

The lysate containing dtSARM1 was pretreated by 100 μM NMN on ice for 10 min, with non-treated dtSARM1 as a control, and incubated with StrepTactin^TM^ XT resin, in the presence of 200 ng/mL Nb-C6, or a CD38 nanobody Nb-1053^25^, overnight at 4℃. The proteins were eluted by 50 mM biotin and analyzed by western blots with anti-His_6_ or home-made anti-SARM1.

### Surface plasmon resonance (SPR)

All SPR analyses were performed with a BIAcore 8K instrument and analyzed by BIAcore Insight evaluation software (Cytiva). To prepare the Strep-capture sensor, CM5 sensors (Cytiva) were coated with Strep-Tactin^TM^ XT protein (IBA) by amine coupling according to the manufacturer’s instructions. Briefly, all channel surfaces were activated by NHS and EDC and immobilized with Strep-Tactin XT protein diluted at 50 μg/mL in 10mM sodium acetate, pH4.5. Excess activated groups were blocked by 1 M ethanolamine-HCl, pH 8.5. A density of Strep-Tactin XT corresponding to 3600- 3800 response units (RU) was achieved.

To measure the binding affinity of Nb-C6 to SARM1, dtSARM, in the lysate of Expi293F, was captured in sample channel to approximately 500 RU, with a blank channel as the reference channel. A series of concentrations of Nb-C6 (0.8 nM-400 nM) in the running buffer, NMN/HBS-EP (100 μM NMN, 10 mM HEPES, 150 mM NaCl, 3 mM EDTA and 0.05% (v/v) Surfactant P20, Cytiva), were injected to both sample and reference channels for 360 s to allow association, followed by 540s of the blank HBS-EP for dissociation. The equilibrium dissociation constant (*K*_D_) value was calculated from kinetics using 1:1 binding model on the evaluation software (Cytiva).

To correlate the Nb-C6 binding with the concentrations of NAD and NMN, the following program was conducted. After dtSARM1 was captured on the SPR sensor, HBS-EP with 2 mM NAD and 200 nM Nb-C6 was injected for 5 min to saturate the nonspecific binding and establish a baseline. HBS-EP containing 200 nM Nb-C6 and a fixed concentration of NAD (0.2, 0.4, 0.8, 1.2, 1.6 mM, each for one channel), together with an increasing gradient of NMN (0, 50, 100 μM) was injected for 5 min, and washed with the same buffer without Nb-C6 for 3 min. The Nb-C6 binding was calculated by the accumulated response signals. The binding rate and concentrations of NMN and NAD were fit into a polynomial model by Curve Fitting Tool (MatLab).

### Immunostaining and imaging SARM1 with Nb-C6

HEK293 cells carrying the inducible expression cassette for SARM1 were seeded on coverslips precoated with 0.05 mg/mL poly-L-lysine. After 12-h induction with 1 μg/mL doxycycline and 6-h activation with 100 μM CZ-48, the cellular SARM1 was immunostained by Nb-C6 as the following procedure. The cells were fixed in 1% PFA for 15 min and permeabilized with 0.1% Triton X-100 in PBS for 5 min. After blocking with 10 mg/ml BSA for 1 h, the cells were incubated with C6-mNeonGreen, anti-Tom20 (Santa Cruze), and anti-SARM1 (home-made) at room temperature for 1 h. The fluorescence was developed by incubating with the Alexa Fluor^TM^-conjugated secondary antibodies, and imaged under a confocal microscope (Nikon A1R) with a 60× object, and analyzed by NIS-Elements AR analysis (Nikon). ImageJ and Imaris Bitplane software were used for colocalization analysis of the confocal signals.

### Quantification of cellular cADPR in cells expressing Nb-C6

HEK293 cells carrying the inducible SARM1 expression cassette were transiently transfected with the plasmids encoding the YFP-fusion nanobody, C6-YFP or 1053- YFP, by PEI. Forty-eight hours post-transfection, the cells were lysed with 0.6 M perchloric acid. After centrifugation, the pellets were re-dissolved in 1 M NaOH and quantified by Bradford assay (Quick Start^TM^ Bradford Kit, BIO-RAD), and the supernatants containing cADPR were neutralized with Chloroform/Tri-n-octylamine mixture (volume ratio 3:1) and treated with NADase (home-made). The concentration of cADPR was analyzed by the cycling assay as described previously^35^. The results were presented as pmol cADPR/mg proteins.

### Cryo-EM sample preparation and data acquisition

The pure dtSARM1 and Nb-C6 were mixed in 1:1 molar ratio in 100 mM Tris (pH8.0), 150 mM NaCl, 1 mM EDTA and incubated on ice for 30 min. In total, 2-3 mg/mL proteins were applied to glow-discharged M026-Au300-R20/20 Copper grid (CryoMatrix) and blotted for 3 s at 8 ℃ with 100% humidity and plunge-frozen in VitroBot Mark IV (Thermo Fisher).

Data were collected on Titan Krios electron microscope (Thermo Fisher), operated at 300 kV and equipped with K2 Summit detector (Gatan). The gatan imaging filter was set with a slit width of 40 eV and the nominal magnification was set to 130,000X corresponding to a calibrated pixel size of 1.076 Å. Data were auto-obtained using SerialEM with defocus ranging from 0.8 to 2.5 μm. Total dose of each movie was 50 electrons/Å^2^, fractioned into 39 frames.

### Cryo-EM image processing

4,605 movies obtained using super-resolution mode were subjected to motion correction using the MotionCor2 integrated into relion-3.1.0 with dose-weighting. CTF estimation was performed with CTFFIND4.1 (relion) and 3,609 micrographs were selected manually based on CTF results. Total number of 3,089,111 particles were auto-picked using Laplacian-of-Gaussian method. After several rounds of 2D and 3D classification, 2,384,641 particles were selected and exported into cisTEM-1.0.0. The particles were subjected to 1 round of 2D classification and 3 rounds of 3D classification with C8 symmetry. A final particle stack composing 208,299 particles was selected based on refinement result showing a final reconstruction map exhibited an overall resolution of 3.0Å with relatively poor density around the ARM and TIR domain. To improve local resolution around the ARM and TIR domain, the particle stack was exported into relion-3.1.0 and expanded with C8 symmetry using relion_particle_symmetry_expand module. Expanded particles were applied to additional 3D classifications in relion-3.1.0, the classes with higher resolution around the outer region of ARM domain and TIR domain were selected and combined after each round of 3D classification. After 3 rounds of 3D classification, 304,208 particles were selected and subtracted with mask that left only “one layer” of the “two-layer” SARM1-C6 complex, leading to a single-layer hexadecimer composed of Nb-C6 and SARM1. The final construction of these subtracted particles was carried out with C8 symmetry, resulting in a final map of >5Å resolution around most part of ARM and TIR domain. Local resolution was estimated by Resmap, map sharpening was performed with lowest resolution for auto-B fit set as 3.5Å.

### Model building and structure refinement

The SARM1^NMN^/Nb-C6 model was built based on the structure of inhibited state SARM1. Our previous reported model of SARM1 binding to inhibitor dHNN (PDB 7DJT) was used as the starting models. SAM1-SAM2 region (403-549), ARM domain (60-402) and TIR domain (561-701) was separately docked into refined map using Dock in Map module of Phenix 1.16. The docked structure was further modified manually in COOT 0.9. Structure model of Nb-C6 was built based on another Nb with similar overall structure (PDB 5F1K) and modified in COOT 0.9. The modified model was further refined by Real-space refinement module of Phenix 1.16 with C8 symmetry maintained during the whole process. Refined models were manually adjusted according to the density and final model was verified in Phenix. The final coordinates and maps were deposited with PDB code of 7XCX and EMDB code of EMD-33129, respectively.

### Analysis of the activity of SARM1 mutants

The mutants of SARM1, including W103A, R110A, K193M, D317A, D317R, E189Q, L257C, S319F, S319Y, Q320A, Q320Y, and F476C were produced by introducing the mutations into the full-length DNA sequence of SARM1 by overlap extension PCR. The resulted mutant genes were subcloned to pENTR1A-GFP-N2 and transferred into the lentivector, pInducer20 (#44012, Addgene) by Gateway system. HEK293 cell line stably expressing SARM1 mutants were established by lentiviral infection and puromycin selection. After induction by 1 μg/mL doxycycline for 24 h, the mutants were extracted with PBS containing 4 mM digitonin and protease inhibitors (Roche).

The activities of mutants were analyzed with PC6 assay with or without pre-activation by 100 µM NMN or acid, as described previously^23^. For the acid treatment, the mutants were incubated with 0.2 M CH_3_COONa (pH 4.3) for 30 s and neutralized with 2 M Tris-HCl (pH8.0). The kinetics of fluorescence (Ex390/Em520) production catalyzed by SARM1 was measured after adding the reaction reagents containing 50 μM PC6 and 100 μM NAD in 50 mM Tris-HCl (pH 7.5) on a Synergy H1 Hybird Reader (BioTek). The initial rate (RFU/min) of the reactions were calculated to quantify the activities of SARM1. The same amounts of lysates were used for western blots.

### Hydrogen-deuterium exchange mass spectrometry (HDX-MS)

HDX was performed by pre-incubating 20µM dtSARM1 with 2 mM NAD or 200 µM NMN at 25℃ for 10 min. After equilibration H/D exchange was carried out by 20-fold dilution into a D_2_O buffer (25 mM HEPES pD7.0, 150 mM NaCl) at 28℃ for 10 s, 30 s, 1 min, 5 min or 30 min (Final 95% D_2_O). The H/D exchange was quenched with additional ice-cold quenching buffer (400 mM KH_2_PO_4_/H_3_PO_4_, pH 2.2, 2.85 M GuHCl) and injected immediately into HPLC (UltiMate 3000, Thermo Fisher) with in-line peptic digestion and desalting steps. Desalted peptides were eluted and directly analyzed by an Orbitrap QExactive HFX mass spectrometer (Thermo Fisher). Non-deuterated samples peptide identification was performed via tandem MS/MS experiments and analyzed by Proteome Discoverer 2.5 (Thermo Fisher). Mass analysis of the peptide centroids was carried by HDExaminer v3.1.2 (Sierra Analytics, Modesto, CA), followed by manual verification for each peptide. No corrections for back exchange that occurs during digestion and LC separation were applied. For peptides exhibiting a bimodal distribution all intensity values belonging to one incubation time in D_2_O were analyzed by fitting two Gaussian peaks with different means and areas but with similar width into the spectra, as described earlier^36^. Next, the fitted peak area parameter was used to calculate the relative amount of open state by taking the ratio of high and the sum of high and low mass subpopulation.

### Crosslinking mass spectrometry (XL-MS)

Crosslinking sample preparation was started with pre-incubating dtSARM1 (in 20 mM HEPES pH8.4) with 2 mM NAD or 200 µM NMN on ice for 10min. Ligand binding SARM1 was incubated with DSBU crosslinker (Thermo Fisher) to a final concentration of 200 µM for 30 min at 25℃, and quenched with 10 mM ammonium bicarbonate. The proteins were denatured with 10 mM DTT and 8 M urea for 60 min, then alkylated by 50 mM chloroacetamide (CAA) for 30 min in dark. Then the protein was digested by trypsin (trypsin: protein = 1:20, w/w) overnight at 37℃. The desalted peptides were separated by EASY-nLC 1200 and analyzed by Orbitrap Eclipse mass spectrometer (Thermo Fisher) using sceHCD-MS2 and CID–MS2–MS3–EThcD–MS2 fragmentation approach essentially as described earlier^37–39^. Proteome Discoverer 2.5 (Thermo Fisher) with XlinkX software 2.5 was used to analyze and identify the crosslinking peptides and the parameters were as follows: maximum of three missed cleavage sites for trypsin per peptide; cysteine carbamidomethylation as fixed modification and methionine oxidation as dynamic modification. Search results were filtered by requiring precursor tolerance (±10 ppm) and fragment tolerance (±20 ppm). FDR threshold was set to 1% for all identified crosslinking peptides.

### Data analysis

All experiments contained at least three biological replicates. Data shown in each figure are all means ± SD. The unpaired Student’s *t*-test was used to determine statistical significance of differences between means (\**P*<0.05, \*\**P*<0.01, \*\*\**P*<0.001, \*\*\*\**P*<0.0001). GraphPad Prism 8.0.1 was used for data analysis.

## Acknowledgement

This study was supported by grants from National Science Foundation of China (31871401, 31871403 and 31950410540), Ministry of Science and Technology (Synthetic Biology Special Project of National Key R&D Program, 2019YFA090600; and Foreign Youth Talent Program, QN2021032004L), Shenzhen Science and Technology Innovation Committee (JCYJ20190808163411340, JCYJ20210324125608023, and 2019SHIBS0004) and Shenzhen-Hong Kong Cooperation Zone for Technology and Innovation (HZQB-KCZYB-2020056). We would like to thank the cryo-EM center of Southern University of Science and Technology for cryo-EM data collection and the HPC-Service Station in cryo-EM center of Southern University of Science and Technology for data processing, the advanced mass spectrometry facility (KMS) of Kobilka Institute of Innovative Drug Discovery, the Chinese University of Hong Kong (Shenzhen) for the support, Mengsi SUN (Office of Core Facilities, ShenZhen Bay Laboratory) for the help in SPR analysis, and Sumyu Yang for drawing the illustration diagram.

---

**Fig. S1.**
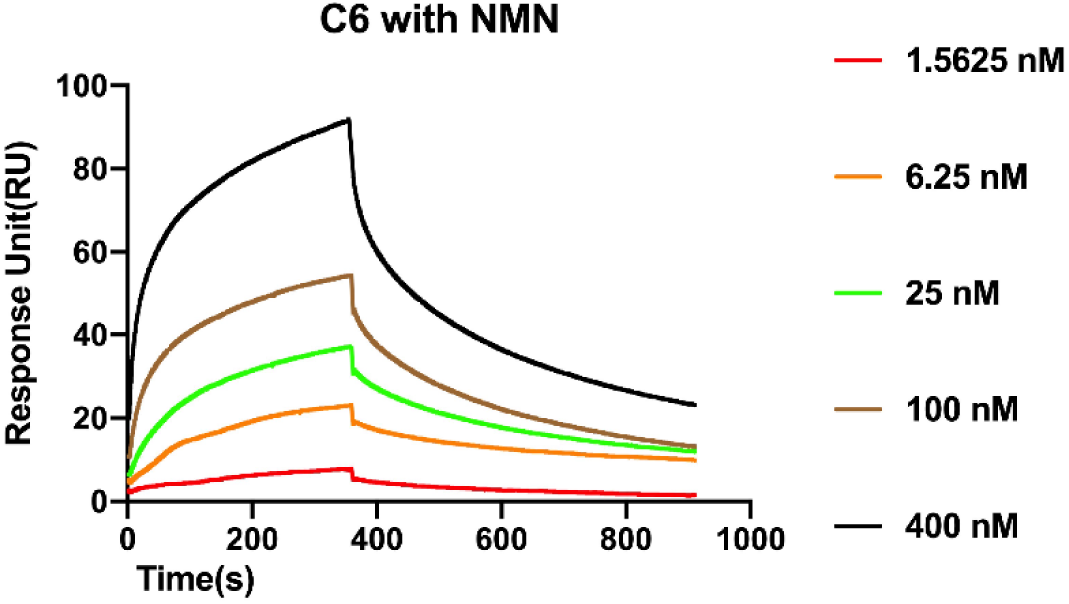
SRP analysis for the affinity between Nb-C6 and SARM1. The recombinant dtSARM1 was immobilized on SPR sensor through the interaction between twin-strep tag and Tactin. A series of concentrations of Nb-C6 (0.8 nM-400 nM) in the running buffer HBS-EP containing 100 μM NMN were injected to both sample and reference channels for 360 s to allow association, followed by 540s of the blank HBS-EP for dissociation. The data were analyzed with the BIAcore Insight evaluation software (Cytiva), giving the association rate constant (*k*_a_) of 2.42×10^5^ M^-1^s^-1^, dissociation rate constant (*k*_d_) of 2.95×10^-3^ s^-1^, and equilibrium constant *K*_D_ of 1.22×10^-8^ M. The data shown is the representative of 5 independent experiments.

**Fig. S2.**
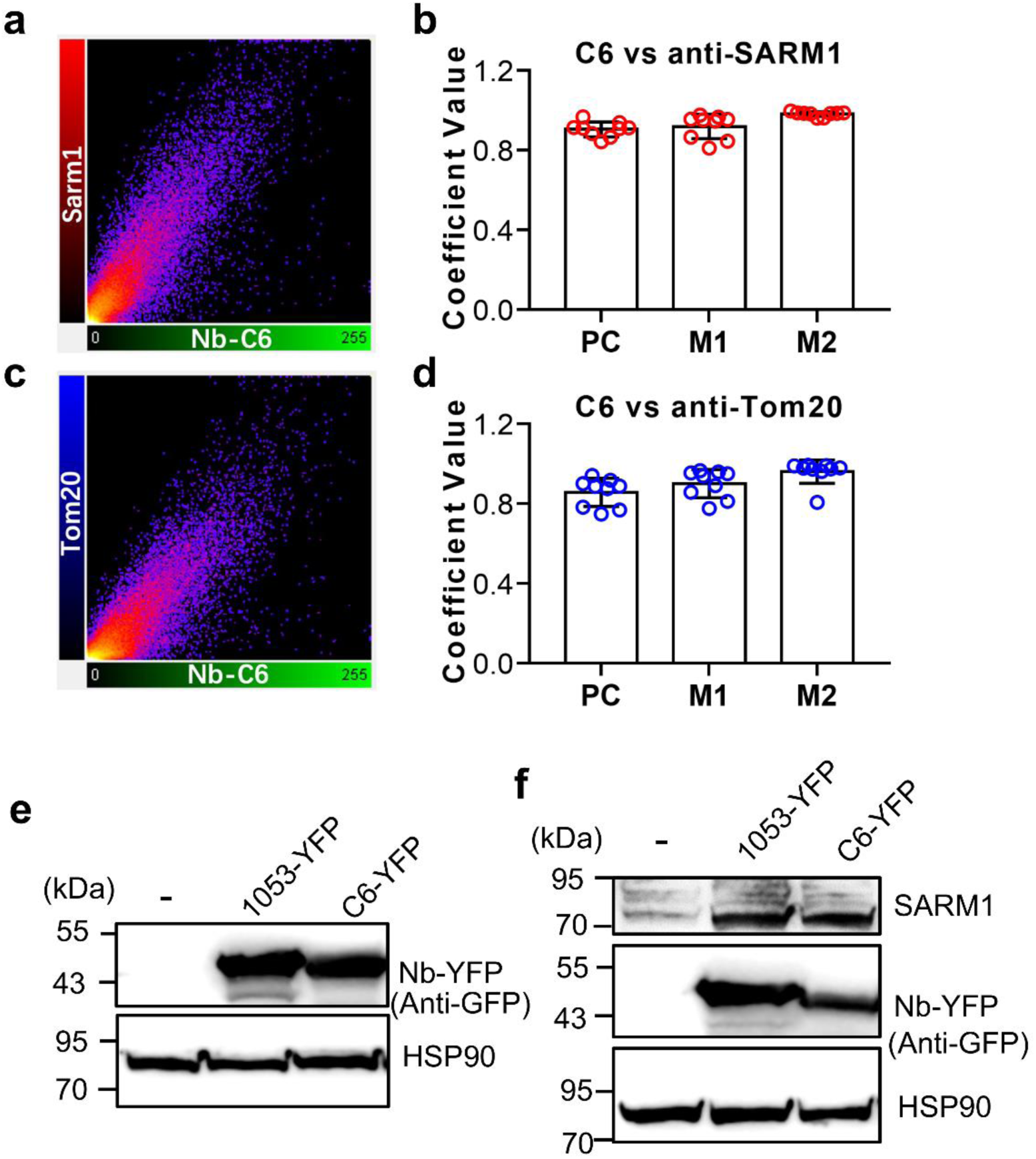
Supplementary data for Fig. 2. (**a-d**) Co-localization analysis of Nb-C6 and anti-SARM1 or anti-Tom20. The pictures are from Fig. 2b (CZ-48 panel). (a and c) Colocalization analysis with Imaris software. Scatter-plot pixels correspond to the images shown in Fig. 2b. Complete colocalization results in a pixel distribution along a straight line whose slope will depend on the fluorescence ratio between the two channels. (b and d) PC, M1, and M2 were analyzed with JACoP. M1 is defined as the fraction of Nb-C6 overlapping anti-SARM1 (red) or anti-Tom20 (blue) signals; M2 is defined conversely. Values are shown as the mean ± SEM of 55 cells. (**e,f**) Protein expression for Fig. 2d and 6e tested by western blots.

**Fig. S3.**
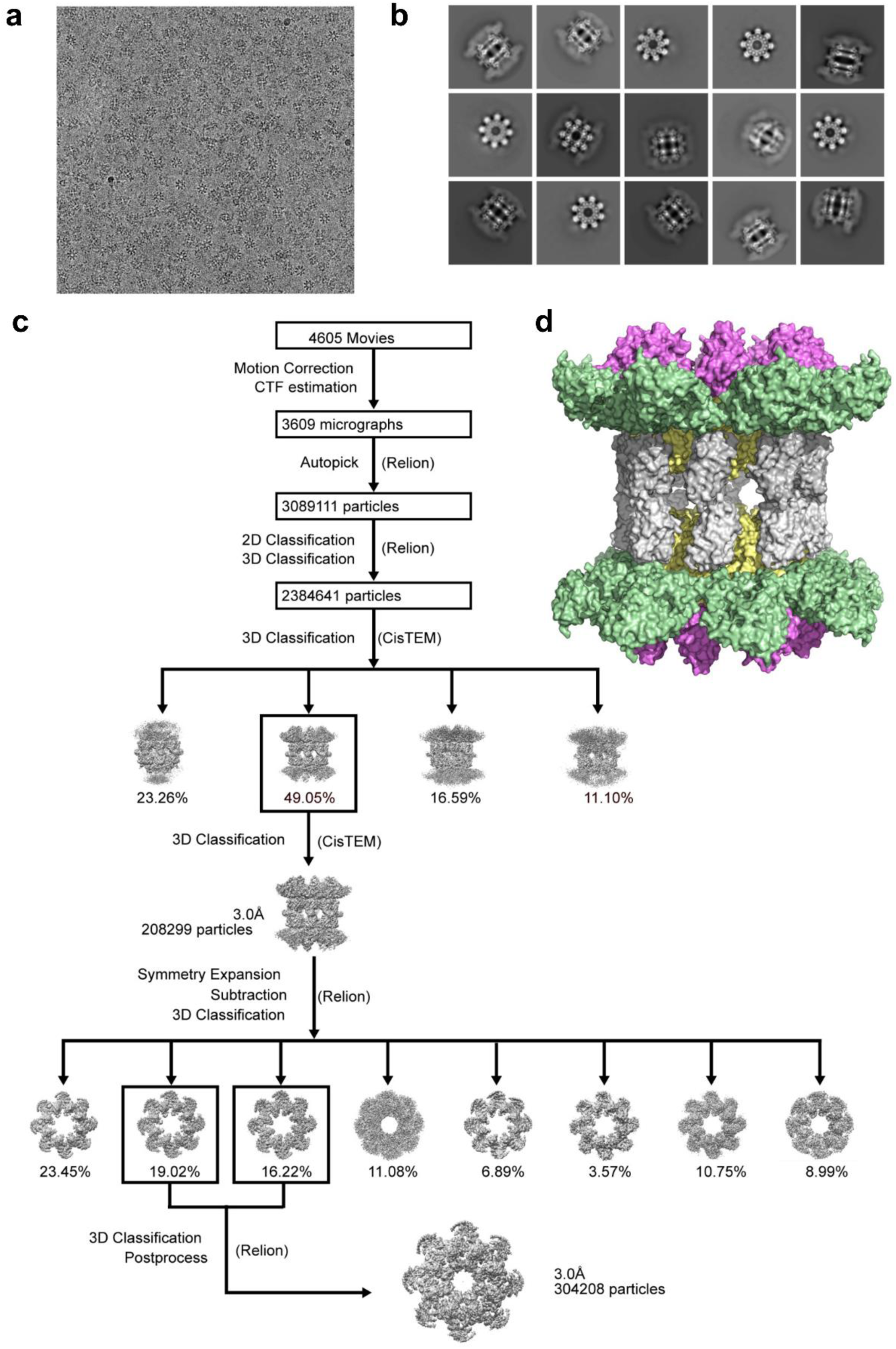
Imaging processing procedure of SARM1^NMN^/Nb-C6 complex. (**a**) A representative cryo-EM micrograph of SARM1^NMN^/Nb-C6 complex. (**b**) Selected 2D class averages. (**c**) Imaging processing workflow of SARM1^NMN^/Nb-C6 complex for refinement and reconstruction of density maps. (**d**) The hexadecameric structure of SARM1^NMN^/Nb-C6.

**Fig. S4.**
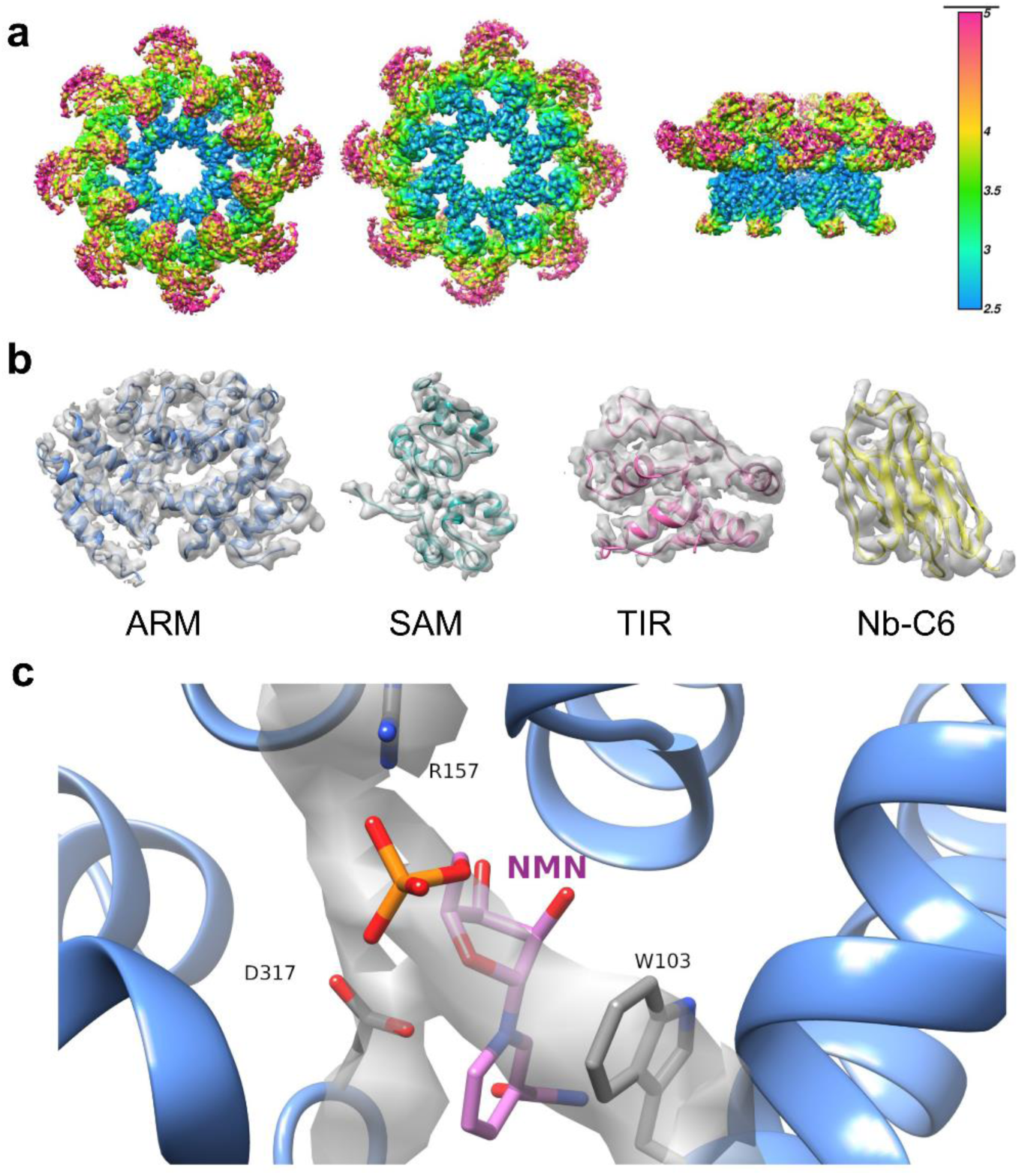
Depiction of density maps of SARM1^NMN^/Nb-C6 complex. (**a**) Depiction of local resolution of the SARM1^NMN^/Nb-C6 octamer in top, bottom and side views from left to right and colored by local resolution generated by Resmap (scale bar shown on the most right panel with blue≤2.5Å, cyan=3Å, green=3.5Å, yellow=4Å and magenta ≥5Å). (**b**) Model fitting of Nb-C6 and each domain of SARM1 into the density maps. (**c**) Local density map for NMN bound in the ARM domain. NMN is shown as stick model (magenta: carbon; red: oxygen; orange: phosphate atoms). Residues W103, R157 and D317 surrounding NMN are also shown as stick models.

**Fig. S5.**
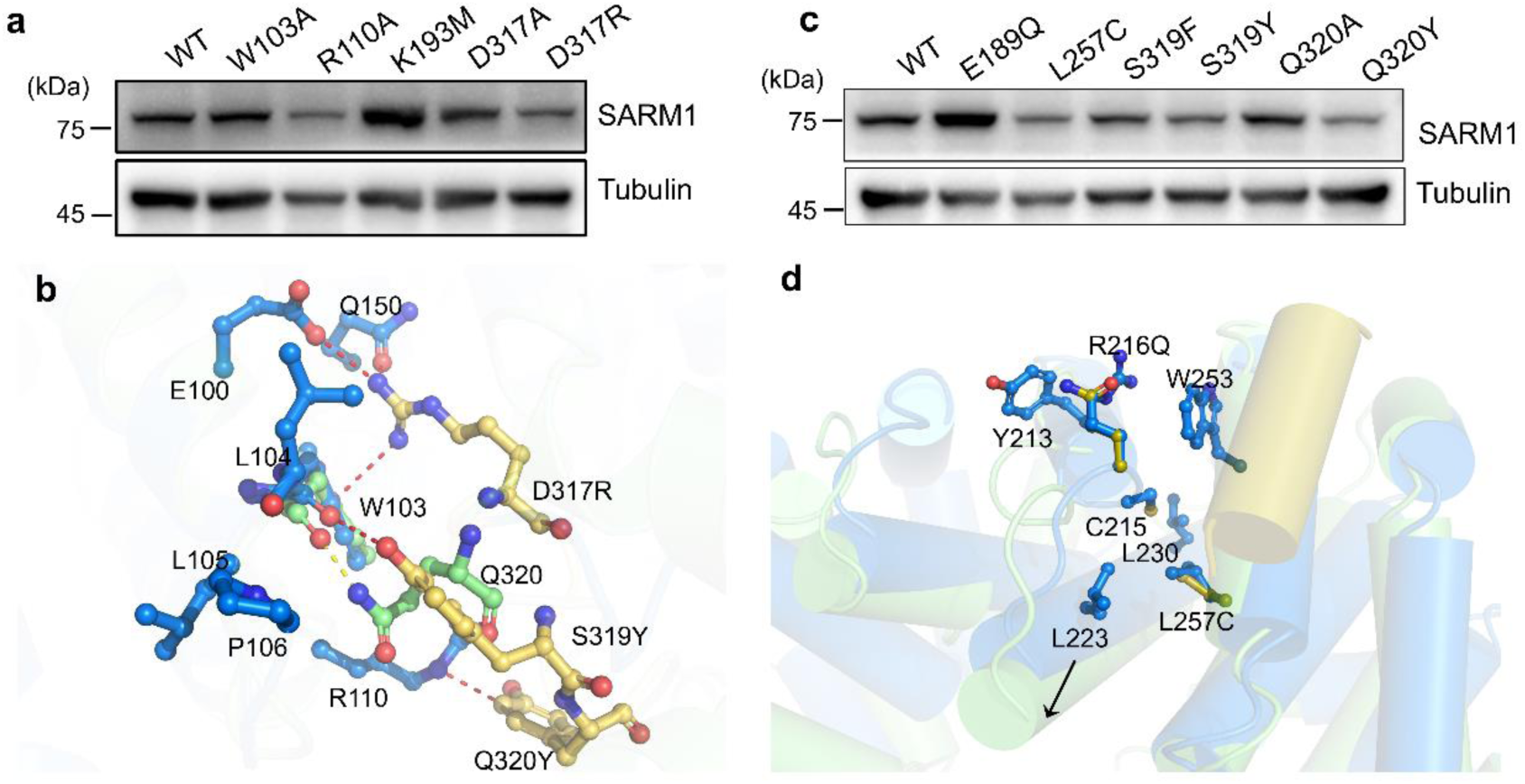
Residues contribute to the NMN-induced conformational change of the ARM domain. (**a,c**) Protein expression for Fig. 4c and 4f tested by western blots. (**b**) Modeling of D317 loop. NMN- and NAD-bound ARM domains (PDB 7ANW) were superimposed via helix 95-105. The color codes are the same as panels **Fig. 4a-b**. D317R, S319Y and Q320Y in ARM^NAD^ were modelled to inspect the interaction with surrounding residues. D317R forms salt bridge with the side chains of residues E100 and Q150 and also positive charge-pi interaction with residue W103, which would stabilize the inactive conformation of the ARM domain. D317E mutant should have similar interaction with H150. S319Y would form hydrophobic interaction with residues L104, L105 and P106 and also H-bond with main chain carboxylate of W103. S319F mutant should form similar interactions to that of S319Y. Q320Y mutant might form charge-pi interaction with residue R110. The D317R, S319Y/F and Q320Y mutants would stabilize the inactive conformation of ARM domain, preventing the inward movement of D317 loop and consistent with the enzymatic results. In ARM^NMN^, the side chain carboxlamine of Q320 forms H-bond with the main chain carboxylate of residue W104. Q320A mutant should diminish the H-bond and weaken the NMN-activation, consistent with the enzymatic result. (**d**) Modelling of mutations at R216 and L257. The ARM-TIR domains of NMN-bound (shown in green) and NAD-bound SARM1 (shown in blue) were superimposed via the helix from AA 575-585 (shown in yellow). The shift of helix from AA 221-236 was indicated by a black arrow. R216Q and L257C mutants were modelled and shown in gold. In SARM1^NAD^, R216 might form pi-Arg-pi interaction with Y213 and W253 and even salt-bridge with residue E689 in the TIR domain. R216Q mutation diminished all these interactions. L257C mutation weakened the hydrophobic interaction with residues L223 and L230 in the helix (221-236), and might even form disulfide bond with C215, facilitating the inward shift of the helix (221-236) induced by NMN. The enzymatic measurements of R216Q and L257C mutants are consistent with the modeling results.

**Fig. S6.**
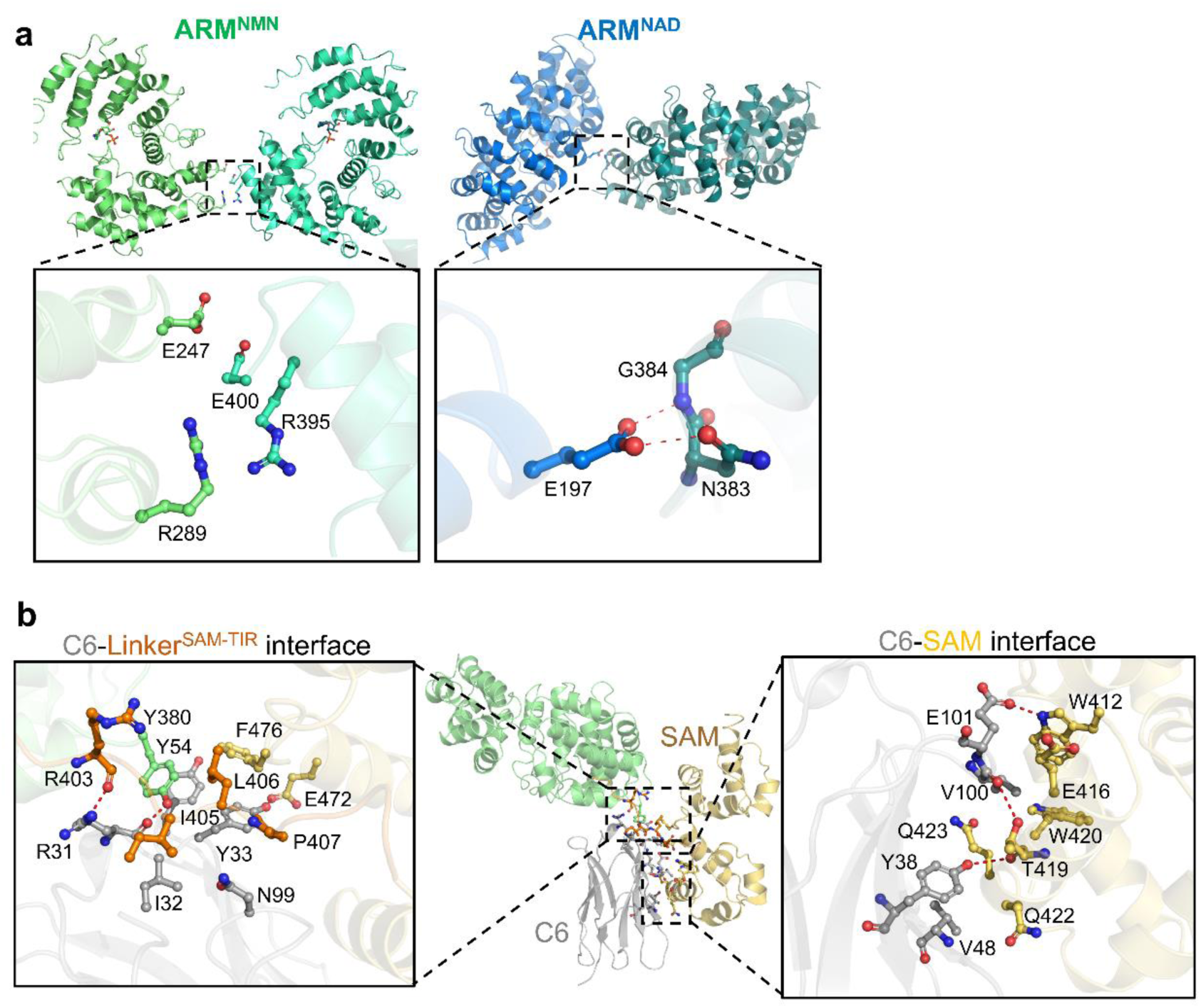
Interfaces between Nb-C6 and SARM1 or ARM and ARM domains. (**a**) Comparison of the ARM-ARM interface in SARM1^NMN^ (left panel) and that in SARM1^NAD^ (PDB 7CM6) (right panel). The adjacent ARM domains in both SARM1 molecules were shown as cartoon models and colored differently. The residues at the interface were shown as ball-and-stick models. (**b**) Interfaces between Nb-C6 and SARM1. Nb-C6 and SARM1 are shown as cartoon models and colored differently for each domain (grey for Nb-C6, green for the ARM domain, yellow for the SAM domain and gold for the ARM-SAM linker). Key interaction residues were shown as ball-and-stick models and labeled. The H-bonds and salt-bridge were shown as red dash lines.

**Fig. S7.**
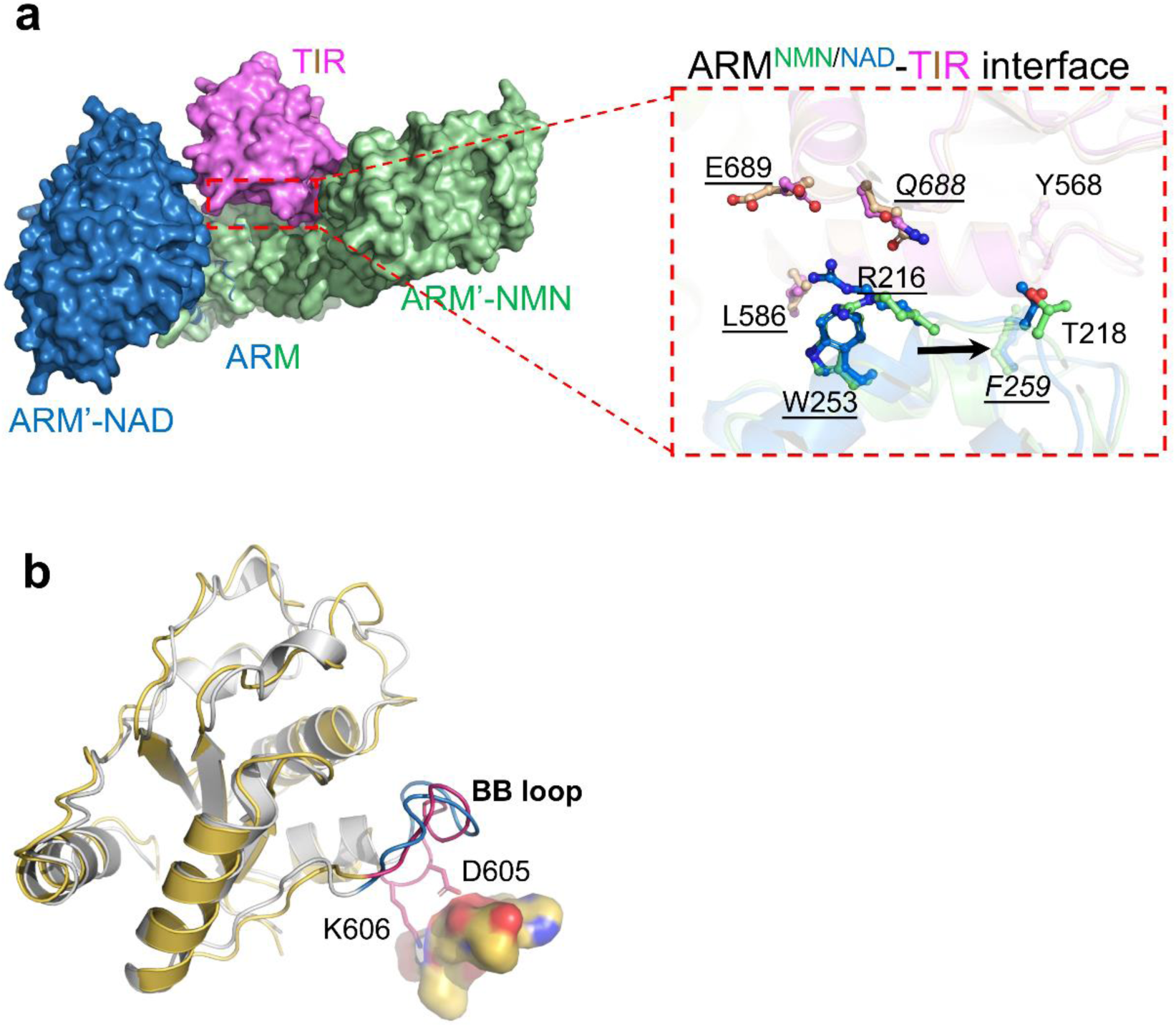
The TIR:ARM interface and TIR conformation are restored in cryo-condition at presence of NAD or NMN. (**a**) Superposition of NMN-bound and NAD-bound SARM1 (PDB 7CM6) via the TIR domains. The TIR domains and two adjacent ARM domains in both structures are shown as surface models and the TIR-ARM interface is boxed and zoomed for detailed view. Some of the residues involved in TIR-ARM interactions are showed in ball-and-stick models. Compared to ARM^NAD^, the helix where R216 and T218 locate moves leftward in ARM^NMN^. (**b**) The superposition of the TIR domains in NMN- (yellow) and NAD-bound (grey, PDB 7ANW) SARM1. The BB loops in NMN-bound and NAD-bound SARM-1 are colored in magenta and blue, respectively. The active site of TIR domain is shown as a magenta ball. Two residues D605 and K606 in the BB loop are shown as ball-and-stick model, which are very close the C-terminus of the SAM domain (E544-P549, shown as surface model) and restrict the opening of the BB loop.

**Fig. S8.**
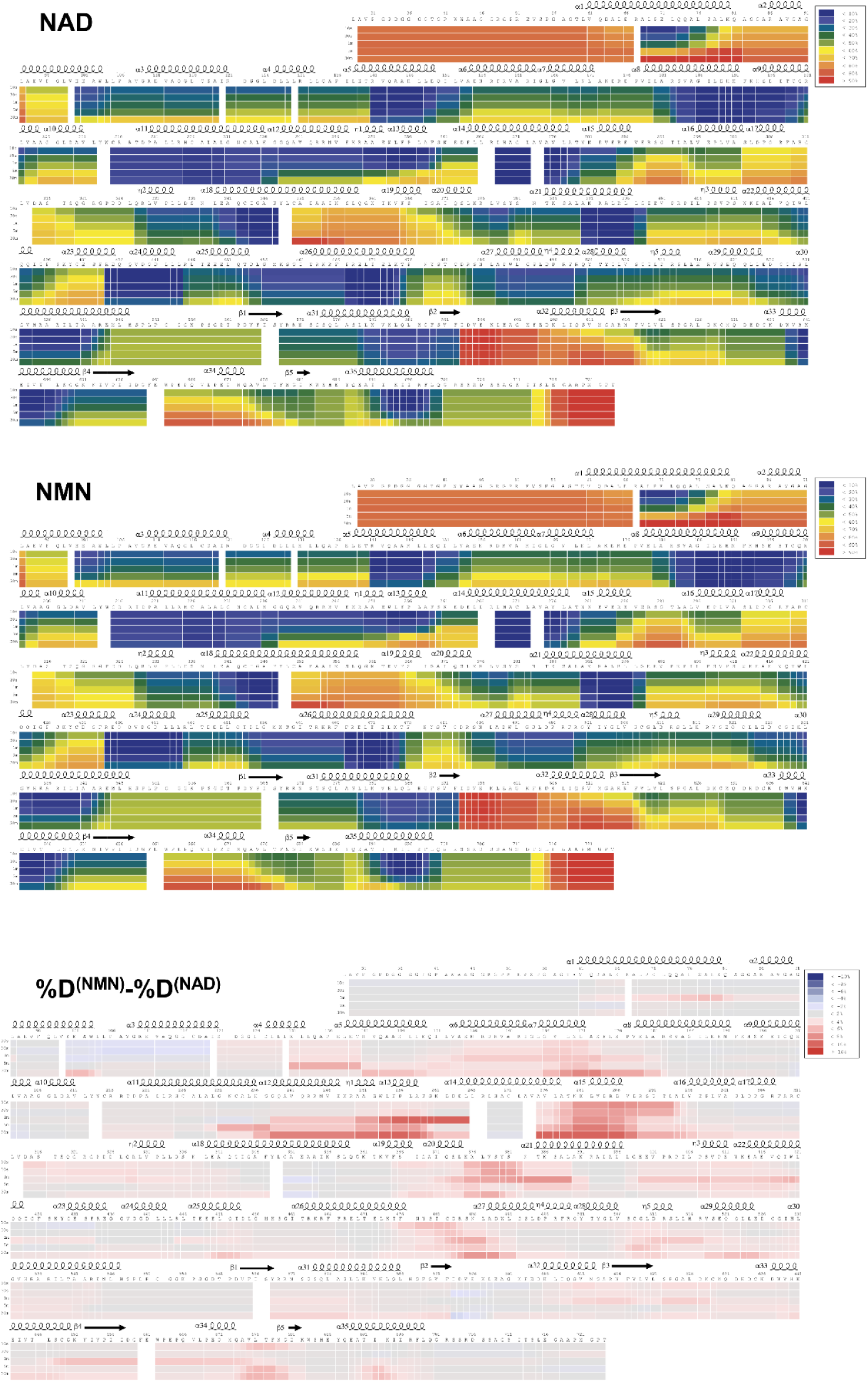
Deuterium uptake data for full-length SARM1 in presence of NAD and NMN. HDX data are shown in heatmap format. Absolute deuterium uptake after 10 s, 30 s, 1 min, 5 min and 30 min is indicated by a color gradient below the protein sequence. Protein secondary structure elements are indicated above the sequence. Bottom panel is showing differences in deuterium uptake between NMN- and NAD-bound SARM1 (%D^(NMN)^-%D^(NAD)^).

**Fig. S9.**
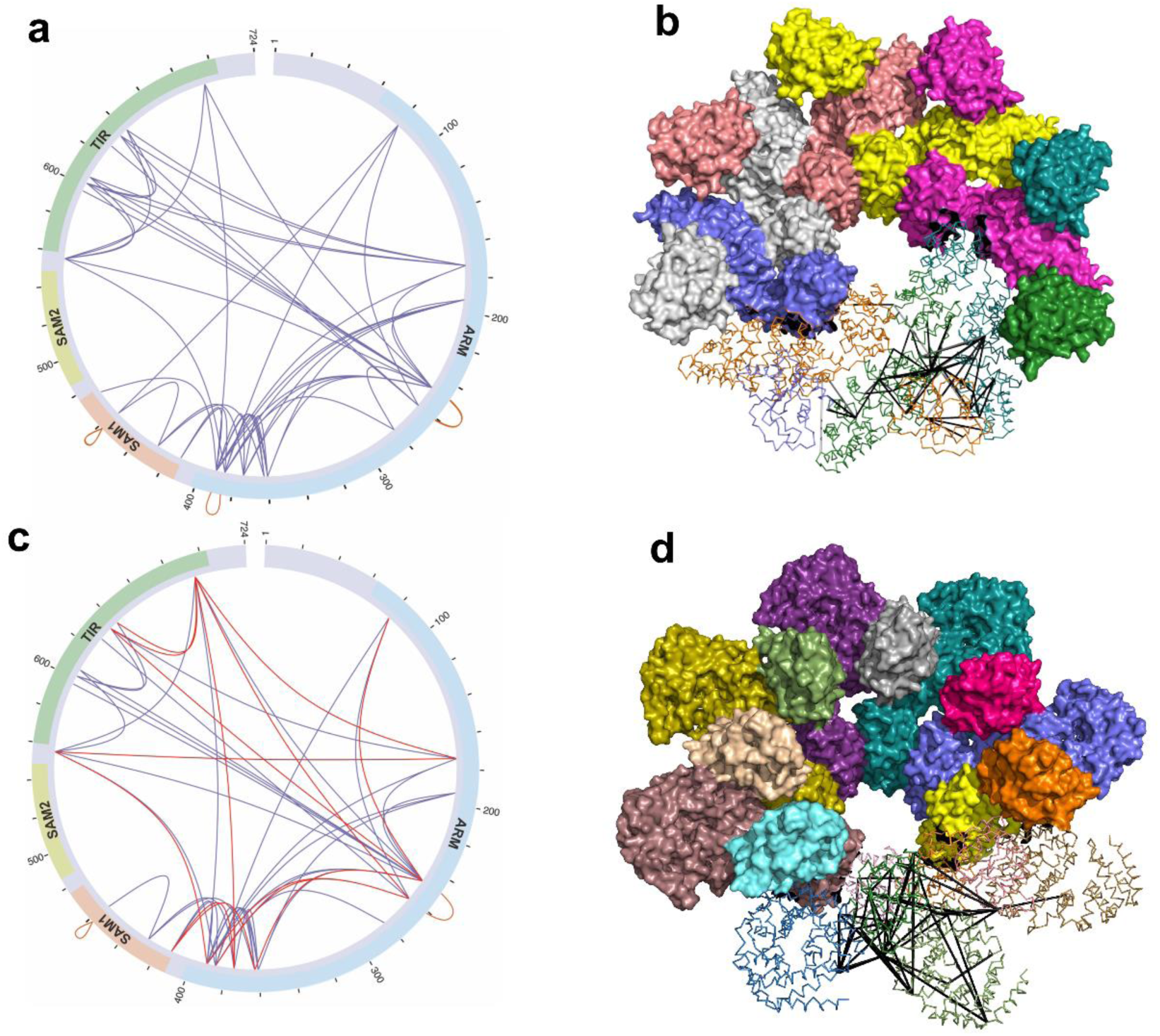
XL-MS analysis of the SARM1 complex in presence of NAD and NMN. Circular plots displaying all the identified XLs for SARM1^NAD^ (**a**) and SARM1^NMN^ (**c**). NMN specific XLs are indicated in red (**c**). Inter- and intramolecular SARM1^NAD^ XLs mapped onto the structures of SARM1^NAD^ complex (PDB 7CM6). (**b**) and SARM1^NMN^(**d**).

**Fig. S10.**
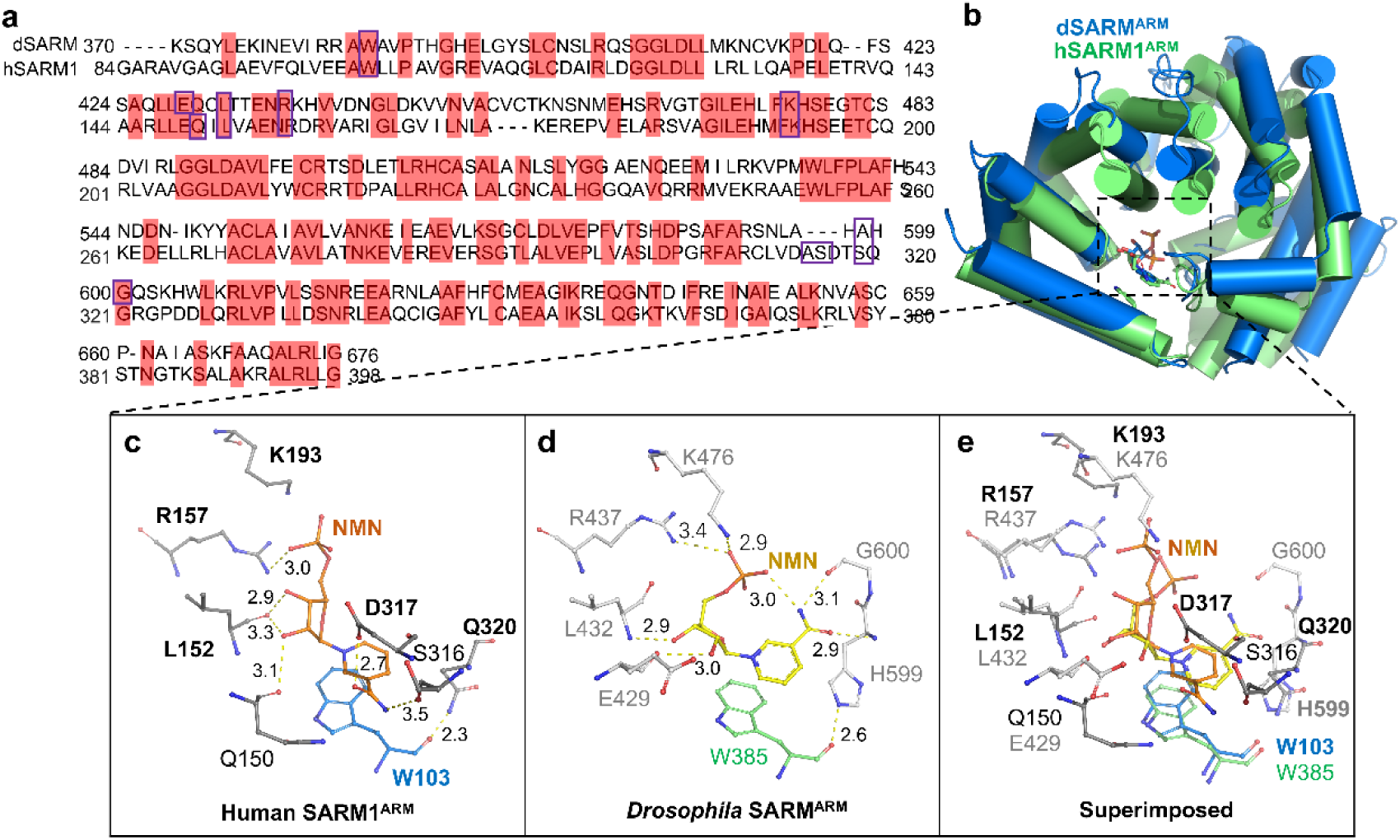
Comparison between the NMN-bound ARM domains of human SARM1 (hARM^NMN^) and *Drosophila* SARM1 (dARM^NMN^, PDB 7LCZ). (**a**) Sequence alignment of the ARM domains in human SARM1 (UniProtKB: Q6SZW1) and *Drosophila* dSARM (UniProtKB: Q6IDD9). Identical residues were highlighted in red and residues interacting with NMN were boxed. The ARM domain of the *Drosophila* SARM1, dARM, which shares 45% of sequence identity with the ARM domain of human SARM1 (hARM). (**b**) Superposition of ARM^NMN^ in human (green) and *Drosophila* (blue) SARM1. NMN was shown as ball-and-stick models. They both show in a helix-composed upside down ω-like shape. (**c**-**e**) Zoomed view of the interaction between NMN and surrounding residues. NMN molecules were colored in gold and yellow, respectively. Surrounding residues were shown as ball-and-stick models with grey carbon atoms. The NMN molecule in dARM orients similarly to that in hARM except the nicotinamide moiety which flips 180° and points downward forming H-bonds with the main-chain amine and carboxyl groups of A598 and G600 (corresponding to S319 and G321 in hARM), respectively. Majority of the NMN-interacting residues are conserved in dSARM, except several such as S316 and D317. Although ARM^NMN^ from *Drosophila* SARM1 (dARM^NMN^) also proceeds conformational changes, the structure is closer to hARM^NAD^ (RMSD of ∼2.1 Å) than to hARM^NMN^ (RMSD of ∼2.4 Å).

